# Nets-within-nets for modeling emergent patterns in ontogenetic processes

**DOI:** 10.1101/2021.02.15.430983

**Authors:** Roberta Bardini, Alfredo Benso, Gianfranco Politano, Stefano Di Carlo

## Abstract

Ontogenesis is the development of an organism from its earliest stage to maturity, including homeostasis maintenance throughout adulthood despite environmental perturbations. Almost all cells of a multicellular organism share the same genomic information. Nevertheless, phenotypic diversity and complex supra-cellular architectures emerge at every level, starting from tissues and organs. This is possible thanks to a robust and dynamic interplay of regulative mechanisms.

To study ontogenesis, it is necessary to consider different levels of regulation, both genetic and epigenetic. Each cell undergoes a specific path across a landscape of possible regulative states affecting both its structure and its functions during development. This paper proposes using the Nets-Within-Nets formalism, which combines Petri Nets’ simplicity with the capability to represent and simulate the interplay between different layers of regulation connected by non-trivial and context-dependent hierarchical relations.

In particular, this work introduces a modeling strategy based on Nets-Within-Nets that can model several critical processes involved in ontogenesis. Moreover, it presents a case study focusing on the first phase of Vulval Precursor Cells specification in *C. Elegans*. The case study shows that the proposed model can simulate the emergent morphogenetic pattern corresponding to the observed developmental outcome of that phase, in both the physiological case and different mutations. The model presented in the results section is available online at https://github.com/sysbio-polito/NWN_CElegans_VPC_model/

## 1. Introduction

Ontogenesis is one of the key concepts at the base of developmental biology [68], defined as “[…] *the development of a single individual, or a system within the individual, from the fertilized egg to maturation and death*” [62]. Ontogenetic processes comprise complex and intertwined mechanisms at different levels, from the embryonic development of the organism as a whole to the differentiation of single cells.

When modeling ontogenesis, a particularly challenging task is to predict the outcome of a developmental process, simulating the formation of emergent morphological and phenotypic patterns from local inter-cellular interactions. A model serving this purpose must describe the cellular organization in space and the consecutive temporal stages characterizing the process. Each stage corresponds to a different conformation and regulative set-up involving multiple interacting cells. Since the regulatory states of these cells depend on their neighbors’ relations, such conformations dictate the communication schemes they engage in [31]. This complexity contributes to creating a multi-dimensional and dynamic landscape of inter-dependent regulative states in which cells can fall into [53, 33]. Moreover, one has to consider that physiology and phenotype not only emerge from the system’s subparts. The subparts themselves self-reproduce, determining internal and system-level regulative and structural changes.

To holistically consider the dynamic properties of this hierarchical regulative environment, a computational model must explicitly account for their evolution in space and time. Moreover, it must quantitatively simulate the underlying non-linear regulatory dynamics across different system levels. This requirement imposes the need to model the hierarchical organization of the system. Eventually, biological systems are intrinsically stochastic, i.e., modeling of their complexity must include stochastic behaviors.

Several modeling approaches for Systems Biology based on different formalisms exist, and each of them has strengths that better fit specific aspects of the target problem [7, 24].

Mathematical models have been among the first proposed to represent continuous biological quantities in chemical reactions at the metabolic level and still prevail in Systems Biology [19, 36, 27].

This is particularly true for the sub-molecular scale, which relies on natural laws and is typically modeled based on Ordinary Differential Equations (ODEs), and is true also for the level beyond molecules, for example, when focusing on concentration of molecules. In the latter case, a continuous description by ODEs naturally supports a macro-view on the system of interest. The focus is on concentrations and their changes over time. ODEs are suitable for dynamics that occur continuously, evolve in a deterministic manner, progress at a similar speed, and can be easily described by real-valued variables. Extensions can relax these constraints, for example, by introducing delays, stochasticity, or discreteness. All in all, continuous formalisms are suitable to describe the concentration dynamics of homogeneous cellular compartments that involve large numbers of cells [22].

Considering the complexity, biological and biochemical systems are usually non-linear, and ODEs describing these systems are often challenging to solve. Nevertheless, they can be approximated by introducing increasingly efficient numerical analysis integrators that enable handling large systems. Also, parameter estimation in the case of a large network with several parameters may have a high computational cost, and the model’s prediction accuracy may decrease [35].

Stochastic discrete event modeling and simulation, where you can control the granularity of observation (instead of setting the degree of accuracy/error of the calculation), is gaining importance as an alternative modeling approach in Systems Biology. The discrete event view describes the dynamics of a system by distinguishing state changes, i.e., events triggered by the flow of time or the situation [76].

Several biological processes are inherently discrete and qualitative, and many examples could be given [23]. Concentrations do not necessarily change continuously, particularly if the dynamics of a small number of entities, like DNA molecules and plasmids, are modeled [42]. Several interactions (e.g., biological signaling, omics regulation, etc.) can be described by discrete stochastic information related to the involved entities (e.g., molecules, cells) [57]. Also, higher-level phenomena of interest for the description of ontogenetic mechanisms (e.g., cell fate determination) are better described by qualitative information.

Several approaches to modeling and simulation of biological systems often focus on modeling intracellular mechanisms and thus fail to adequately capture supra-cellular spatial elements, limiting their capability to analyze patterns emerging from local interactions between biological entities [7, 6]. Comprising both intracellular and supra-cellular information can broaden the scope of representations. Yet, it poses the challenge of combining heterogeneous information from multiple system levels in a single model. One of the approaches for handling model heterogeneity is the composition of existing models into a more extensive scope [7]. However, this approach raises consistency issues [61] and introduces additional requirements to create multilevel and hybrid models [4].

This paper introduces a computational modeling methodology based on Nets-Within-Nets (NWN), a formalism based on Petri Nets (PN). NWN rely on a single formalism to cover all requirements posed by the modeling of complex ontogenetic processes. The paper does not aim at introducing a new formal extension of the NWN formalisms. NWN are instead used as a powerful instrument to propose a methodology to model complex dynamics, stochastic processes, hierarchical organizations, and spatial structures in biology. NWN support composition and integration processes typical of hybrid models, supporting the integration of heterogeneous sources of knowledge. In other words, NWN leverage the strengths of modeling paradigms from various existing computational tools and methods, with the advantage of maintaining uniformity in formalism usage. This paper explicitly refers to the NWN implementation provided by Renew, an extensible editor and simulation engine for Petri nets [41, 11, 70]. The main advantage of this framework is to add the full power of an object-oriented language (i.e., Java) to the NWN formalism, thus allowing the description of more complex functionalities.

After introducing the NWN formalism, its relations to other PN-based formalisms, and our usage of its capabilities for modeling ontogenetic processes, this paper provides a working example of the proposed approach. This example models a well-characterized and straightforward ontogenetic process: the Vulval Precursor Cells (VPC) specification process in *C. Elegans*. Eventually, the paper provides an appendix proposing a library of NWM modeling basic biological processes that interested readers can use as a starting point to build models of different ontogenetic processes.

## 2. Background

The goal of this section is not to provide a complete and formal review of the PN theory. Interested readers may refer to [59] for a detailed introduction on this topic. This section provides an overview of PN applications in the biological domain and, most importantly, introduces the basic characteristics of the NWN formalism exploited to model ontogenetic processes in the following parts of the paper.

### 2.1. Petri nets definition

Petri Nets (PN) are discrete event system models first introduced in the early 1960s by Carl Adam Petri in his Ph.D. dissertation [56]. PN combine a well-defined mathematical theory with a graphical representation of the dynamic behavior of the systems. This combination is among one of the main reasons for the great success of PN, which have been used to model various kinds of dynamic event-driven systems, including biological systems [12].

A PN is a directed bipartite graph having two types of nodes: *places* representing states or conditions and *transitions* representing events that may produce, consume or take resources from one state to another. Directed arcs can only link places to transitions or transitions to places. Places can contain a discrete number of *tokens*, each one representing a resource unit. The marking of a PN is the token configuration in its places. It can dynamically change when transitions fire. Formally, a PN is a 4-tuple *N* = (*P, T, W, M*_0_), where:

1. *P* = {*p*_1_, *p*_2_, …, *p*_*m*_} is a finite set of places;
2. *T* = {*t*_1_, *t*_2_, …, *t*_*n*_} is a finite set of transitions, *P* ⋃ *T* ≠ ∅, and *P* ⋂ *T* = ∅;
3. *W* : (*S* × *T*) ⋃ (*T* × *S*) ⟶ ℕ assigns to each arc a non-negative integer arc multiplicity (or weight); note that no arc may connect two places or two transitions;
4. *M*_0_ : *P* → *N* is the initial marking, i.e., the initial configuration of tokens. Together with the network architecture, it defines the PN model.

Figure 1 shows an example of the graphical formalism used to represent a simple PN. Specific symbols are assigned to represent places, transitions, arcs, and tokens. Transitions can move tokens from their input to their output places. Firing a transition consumes *W* (*p*_*i*_, *t*) tokens from each input place *p*_*i*_ and produces *W* (*t, p*_*j*_) tokens in each of its output places *p*_*j*_. When not indicated, the weight of an arc is equal to 1. A transition *t* is enabled (it may fire) in a marking *M* if there are enough tokens in its input places for the consumptions to be possible, i.e., if and only if ∀ *p: M*(*p*) ≥ *W*(*p, t*).

**Figure 1:**
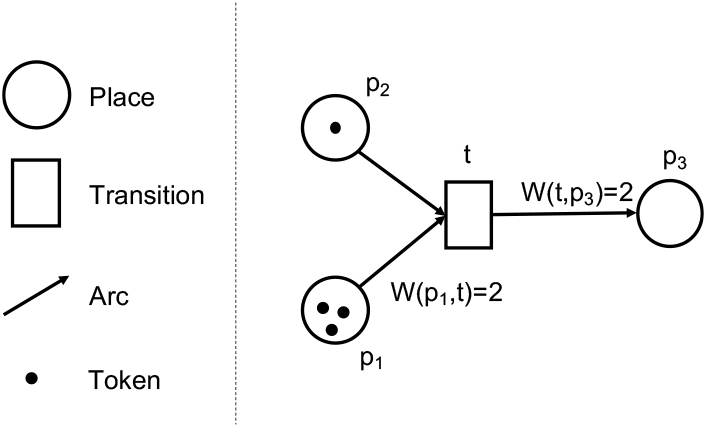
Graphical representation of a simple Petri net. The basic elements that compose a Petri net are places, transitions, arcs and tokens. Transitions are able to move tokens from their input to their output places.

Starting from this basic definition, also referred to as *low-level PN*, several PN variants have been defined. Some of them found application in supporting biological systems modeling to different extents.

### 2.2. Petri nets in biological applications

Simple biological mechanisms such as isolated biochemical reactions require a quantitative representation of kinetic parameters and stoichiometric relations between the involved molecular species. In a low-level PN, places can represent species (substrates and products), while transitions can model reactions. Tokens can express in a discrete way relative quantities of molecules.

Moving from a single reaction to groups of biochemical reactions (e.g., metabolic and gene regulation networks) introduces further requirements such as more complex arhitectures and concurrency [39]. Low-level PN can still support this complexity. In a metabolic network, places can model molecular species and states, while transitions can model enzymatic reactions. Tokens represent discrete quantities of molecules, while the architecture of the network represents how resources flow within a network of competitive or sequential reactions. Similarly, in a regulation network, places can refer to genes and gene products. Transitions can cover transcription, translation, and regulation processes. At the same time, tokens can model elements from the relevant omic pools (e.g., genes, transcripts, proteins) and the regulatory conditions in which each of them can fall.

The phenomena considered so far refer to a single system level (i.e., intracellular mechanisms) and consider a limited temporal scale. Expressing different time scales provides greater expressivity which is often required, for example, to model differential transcriptional rates. To handle different timescales, timed PN [64] introduce a time delay associated with the activation of each transition. Once a transition is enabled, deterministic time delays can occur, ordering the different activation events during the net evolution. The introduction of time delays increases time resolution when including diverse yet intertwined mechanisms in a model.

Stochastic PN [45] extend Timed PN with the use of probabilistic time delays. They can model the stochasticity of a biological system (e.g., gene expression level random fluctuations [29]). The delays become random variables that can depend on the current marking of the net [67].

Time continuous PN have also been proposed in the literature [17]. They can be a valuable tool to model selected biological processes that are not discrete in nature. While these PN can potentially be employed in the methodology presented in this paper, they have not been exploited since not supported by the selected simulation framework.

In general, low-level PN do not scale with system complexity. The lack of scalability limits their use to the modeling of small systems. *High-level* Petri Nets can increase the model’s expressivity, including different system levels and dimensional scales to provide a more systemic view of the biological complexity. High-level PN extend the low-level formalism, supporting multi-level and nested models that properly handle information diversity and complexity [44].

Colored PN (CPN) [34] are the simplest class of high-level PN. They are important since they allow to associate tokens with arbitrarily complex data structures defined as *colors* encoding complex information. Moreover, in CPN, each place and transition can be designed to accept a limited number of colors. In this way, it is possible to separate the identity of resources from their location, modeling the same condition for different categories of resources [50]. CPN lead to non-redundant and more compact representations of the system. This compact representation improves readability and averts modeling errors while preserving the modeling capabilities of low-level PN, which can be generated from CPN models by automatic unfolding [43, 44].

Models considered so far flatten information from different system levels into a single one. To represent a multi-level system structure, hierarchical PN organize system parts and subparts in nested net architectures with explicit hierarchical relations. The nested architecture allows for arbitrarily high resolution when describing mechanisms from different system levels [48]. Nevertheless, like CPN, hierarchical PN stick to a static paradigm: token colors are static data structures, and nets have a fixed model architecture. Resources can change state only by moving from place to place, i.e., changing their position in the net, preserving the information they carry with them unaltered. Also, mobility is devised for tokens, i.e., for resources, but not for other model elements.

Complex biological processes such as ontogenesis challenge the limitations of most high-level PN. Ontogenesis comprises architectural and functional system changes across different phases of the same process. These changes include the movement and generation of new parts and decision-making processes based on previous process stages. For example, an embryonic development process can be considered at different system levels, from the organismal structural rearrangements to local molecular and cellular interactions from which morphological and functional patterns emerge at each developmental stage.

The NWN formalism is an extension of PN in which tokens themselves have the structure of a PN. Tokens specified in this way (net-tokens) evolve dynamically, just like the net holding them (system-net). This hierarchical organization can be reiterated in a boundless way, allowing for open recursion in specifying the system’s hierarchical organization [10]. The idea of tokens being PN goes back to R. Valk [71], and NWN approaches are extensively studied in the PN literature [40, 46, 63, 10]. The Nets-Within-Nets (NWNs) formalism supports all capabilities of other high-level PN and can face the additional modeling requirements posed by ontogenetic processes [5, 3] as will be explained in the next sections. Among the different flavors of the NWN formalisms, this paper adopts the one provided by the Renew simulator [41, 11, 70]. One of the main motivations behind this choice is that it provides a robust framework that allows extending the NWN formalism with the full power of the Java object-oriented programming language (i.e., Java), thus allowing the implementation of more complex functionalities.

### 2.3. An introduction to the Nets-Within-Nets formalism

The NWN formalism introduces the possibility to specify tokens in terms of PN models. Such tokens are called *net-tokens*, or *object nets*, while the net holding them takes the name of *system-net*. This schema can grow recursively to an arbitrarily large number of levels: net-tokens from a level may function as system-nets for net-tokens at a lower one.

Figure 2 shows a simple NWN model used to introduce the basic modeling elements exploited in this paper. The figure uses the NWN notation offered by Renew, the simulation environment this work relies on. This notation is used consistently throughout the paper. The proposed system is composed of a system-net (SN) in which four types of tokens coexist. As in low-level PN, the simplest category of tokens is black tokens, denoted as [] in place 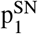. The consumption or creation of tokens by a transition is denoted by inscriptions on its arcs, listing the involved tokens separated by semicolons. Black tokens do not bring specific associated information other than the presence of a unit of a generic resource. Whenever more specific information is required (e.g., a string or an integer value), colored tokens can be used, as shown in place 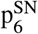. NT1 and NT2 define the architecture of two net-tokens. The inscription N1: new NT1 in transition 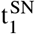 denotes that a new instance N1 of the net NT1 is created,and a reference to this object is then located in place 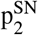. Following the object-oriented paradigm, a reference to the same instance of a net-token can be instantiated in different places. Therefore a net-token can exist at the same time in different places or even in different nets. In Figure 2, the transition 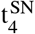 creates two references to the same instance of the same net-token N2 in places 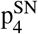 and 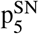. Finally, the bidirectional arc connecting 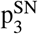 with 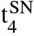 denotes that token N1 is used to fire the transition and then moved back to its original place.

**Figure 2:**
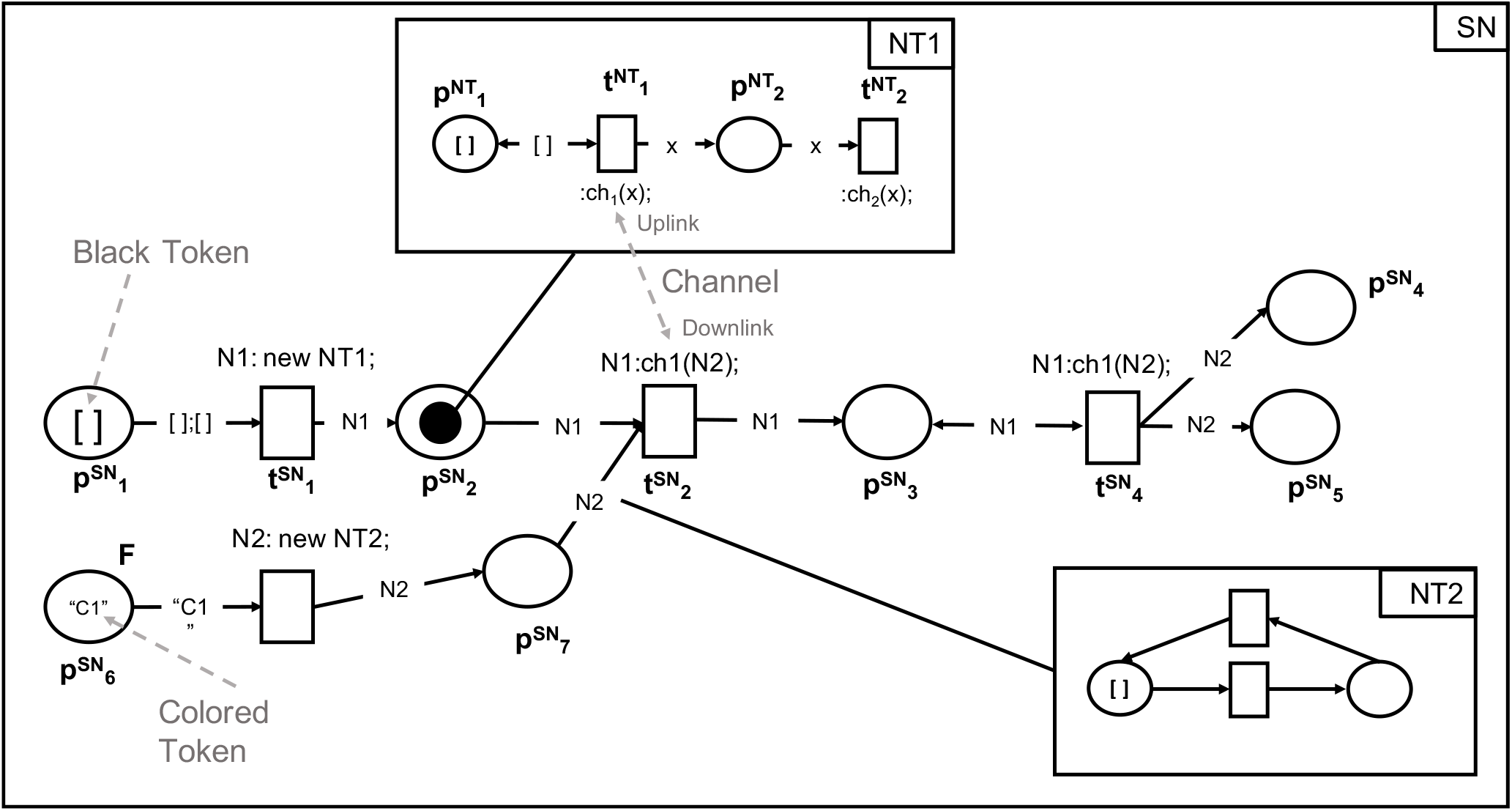
Simple example of the NWN formalism usage. The example represents a system-net (SN) holding the three categories of tokens available in an NWN: (i) black tokens as in low-level PN, (ii) colored tokens as in Colored PN, and (iii) net-tokens that are complex tokens represented again using the PN formalism. Communication channels enable communication between different hierarchical levels of the NWN. Interested readers can refer to [70] for a complete and detailed description of the formalism.

This short overview shows how NWN can express a range of possible scenarios. The following section presents the way these capabilities respond to the modeling requirements posed by complex ontogenetic processes.

## 3. NWN applied to ontogenesis modeling

This section applies the NWN object-oriented paradigm to the problem of modeling ontogenetic processes. It focuses on describing the most relevant and complex biological semantics involved in every ontogenetic process as depicted in Figure 3. After presenting these mechanisms, Appendix A proposes an extensive set of biological processes modeled using NWN. These models represent a library of basic building blocks that can build complex models of generic onto-genetic processes.

**Figure 3:**
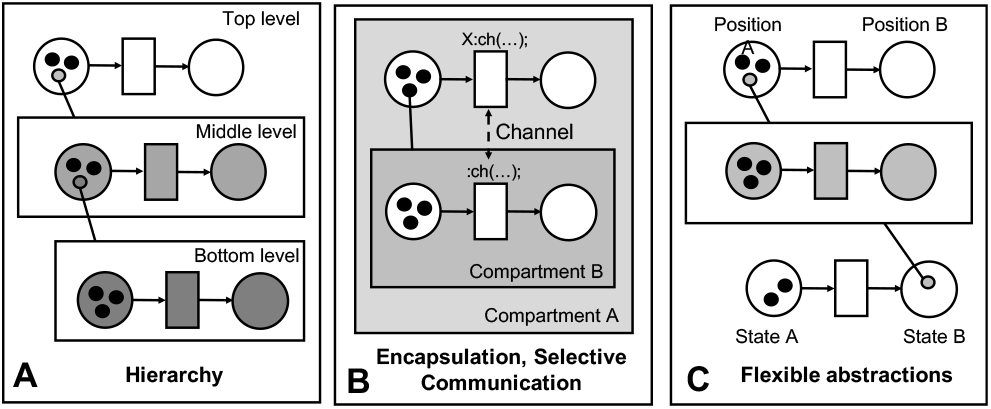
NWN modeling capabilities. NWN models express (A) multiple hierarchical levels through the concept of net-tokens instantiated into a system-net; (B) encapsulation of model parts and selective communication between them through the use of communication channels between different nets; (C) coexistent and intertwined regulatory layers with different annotations thanks to the possibility for a net-token to be referenced in several system-nets.

In our approach, the net-tokens represent differentiating biological cells described through their internal regulative networks. The system-net instead represents the landscape of external factors that affect the cells’ functioning. The system-net includes the microenvironment (e.g., environmental factors, spatial organization, and relative cell positions) and the relevant developmental phases. The presentation of the proposed modeling strategy resorts to a simple abstract example, depicted in Figure 4. The target biological system is composed of a Petri dish divided into four subspaces (A) where two cells of the same type and able to exchange and react to signals (B) coexist. In the first stage of the process (A), cell NT1 moves in a subspace closer to cell NT2 (C), and the two cells start exchanging biological signals mediated by mutual activation of signaling mechanisms (D). As a result of this interaction, the process enters a new stage (E) where the cells change their state (F).

**Figure 4:**
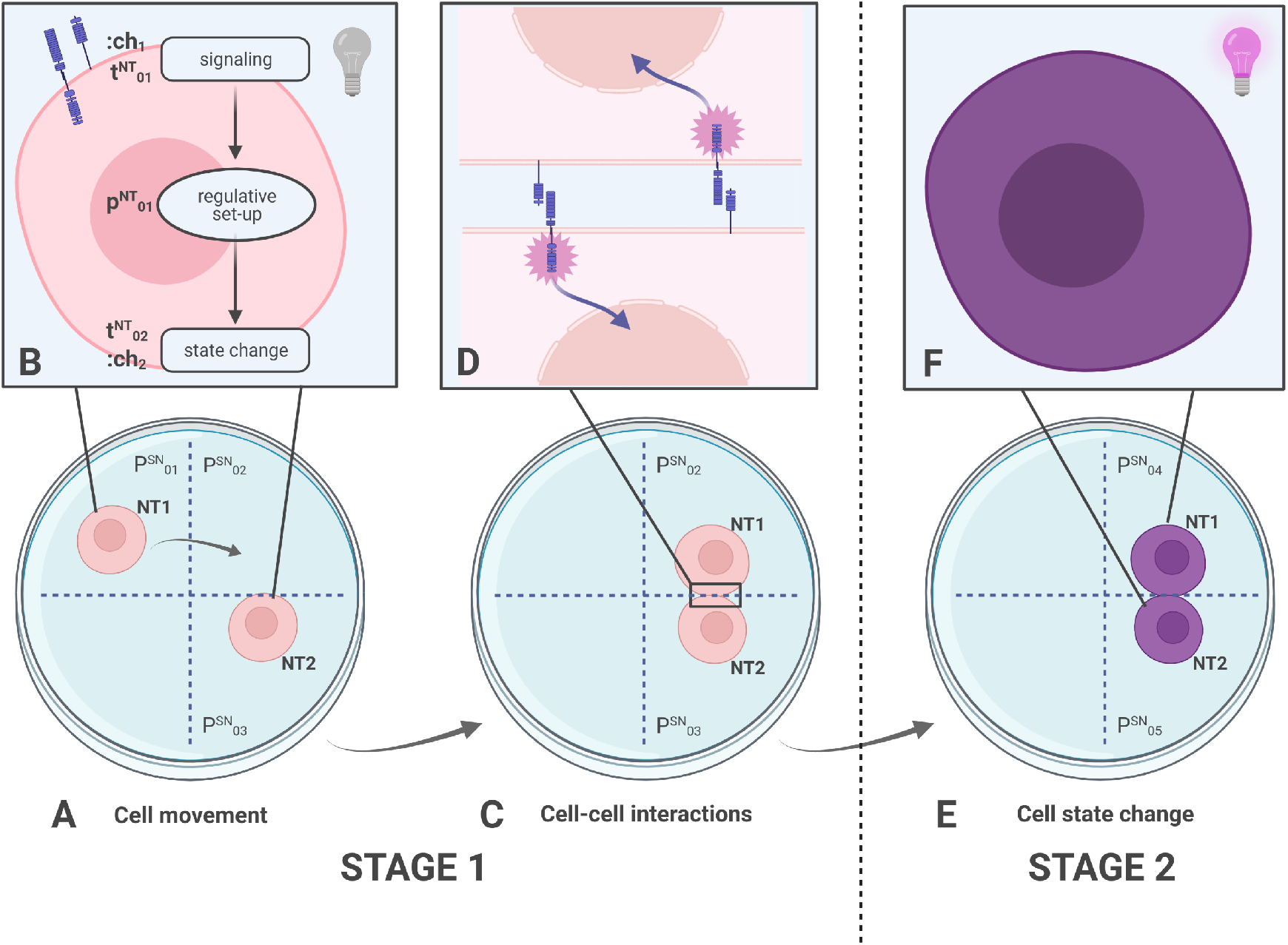
A simple abstract example of an ontogenetic process. The system involves (A) the movement of a cell (NT1) between different subspaces of a Petri dish, considered in reason of (B) the signals it exchanges when in contact with other cells, the regulative set-up it acquires after exchanging signals, and the following changes in cell state. (C) The interactions between cells in spatial proximity, which are mediated by (D) mutual activation of signaling mechanisms, lead to (E) a change in the cells state depicted here with a change of color of the cell (F) that moves them to the second stage of the ontogenetic process (E).

Figure 5 shows a simple NWN modeling the mechanisms presented in Figure 4. NT1 and NT2 model the two cells as instances of the net-token (NT). The system-net (SN) models the spatial organization of the Petri dish and the process organization in the two considered stages.

**Figure 5:**
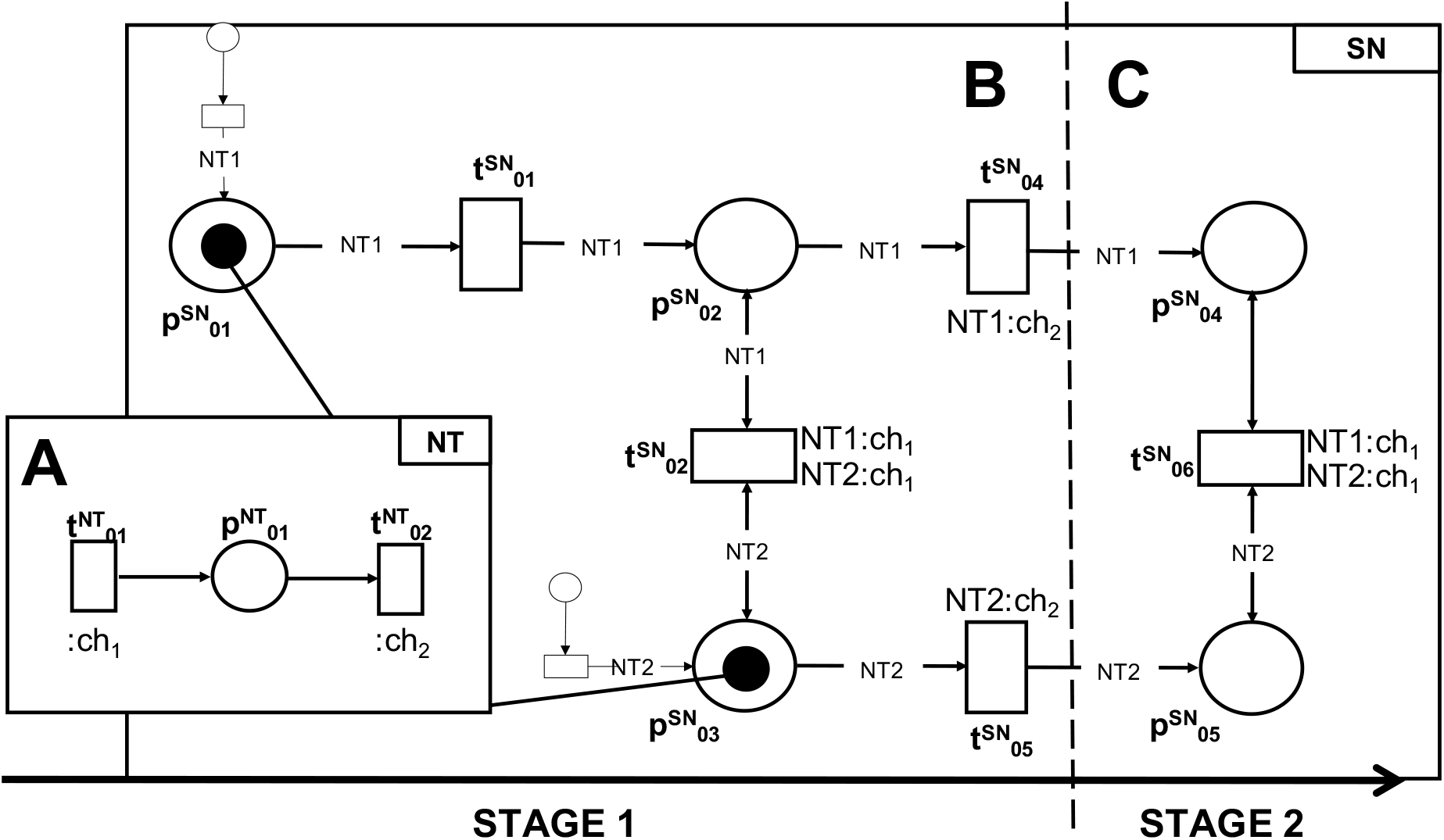
A simple NWN modeling the biological process presented in Figure 4. The system-net (SN) uses places to represent locations that various actors occupy. Places 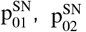, and 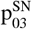 represent the three subspaces of the Petri dish in which cells represented by instances of the net-token NT can live during stage 1 of the ontogenetic process. Similarly, 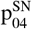 and 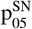 are the two locations of interest during stage 2. The movement of cells from one location to another one is modeled using transitions. Interactions between cells existing at different positions are modeled using communication channels. Finally, the net-token represents the internal behavior of the involved cells.

### 3.1. Spatiality and mobility

NWN models can explicitly represent spatiality, using places to represent actual locations that various actors occupy. For example, in Figure 5, places 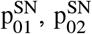 and 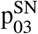 represent the three subspaces of the Petri dish inwhich cells (net-tokens) can live during stage 1 of the ontogenetic process. Similarly, 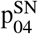 and 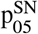 are the two locations of interest during stage 2.

Interactions between the actors existing at different positions can model proximity-enabled communication between neighbor cells involving communication channels. For example, transitions 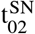 and 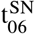 express proximity between 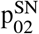 and 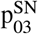, and 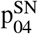 and, 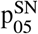 respectively. When these transitions fire, they involve the net-tokens from the connected places, creating mutual interactions. This mechanism can represent juxtacrine interactions between neighbor cells in a biological context, which occur only when the cells are in close spatial proximity.

The movement of net-tokens across system-net places having spatial semantics models mobility of biological entities. For example, in Figure 5, transition 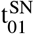 can move net-token NT1 from the place 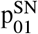, representing a position in space, to the place 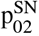, representing a different position. In a biological system, this corresponds to a cell’s movement from a location to another. In general, biological actors, including cells, are capable of active movement. Developmental processes often comprise cell migration phases across different microenvironments.

Net-tokens that move from one system place to another one change the set of interactions they can engage. When involving places with a spatial connotation, this mechanism models the regulatory action of the spatial context for biological entities. The shift from a biological microenvironment to another corresponds to a regulatory change since it comes with potential engagement in interactions with different actors.

### 3.2. Semi-permeability of biological compartments

The object-oriented paradigm implemented by the NWN formalism is powerful to express *encapsulation* and *selective communication* mechanisms. These features, coupled with the ability to handle spatial information, make it easy to describe *compartmentalization* and *semi-permeability* of membranes between biological compartments.

In our modeling approach, the net-tokens describe the inner functioning of cells intended as biological compartments. For example, net-tokens NT1 and NT2 in the system-net of Figure 5 are two instances of the same net class, depicted in Figure 5-A. The system-net can interact with the net-tokens dynamics of both instances through the synchronous channels ch_1_ (transition 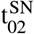 and 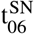 in Figure 5) and ch_2_ (transitions 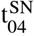 and 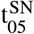 inFigure 5). Biological compartments are semi-permeable: communication mechanisms between the net-tokens and the system-net represent the selective permeabilities of biological membranes, including the mechanisms regulating them.

### 3.3. Inter-cellular communication

In developing multicellular organisms, each cell behaves independently from the others yet can exchange signals and resources. These interactions create the complex and highly dynamic regulative landscape in which cells live and move.

Paracrine and juxtacrine signaling depicted in Figure 6 are among the most relevant intercellular communication mechanisms underlying ontogenetic processes. In biology, paracrine signaling is a form of cell signaling in which a cell produces a signal and sends it to the extracellular environment, affecting nearby cells. Juxtacrine signaling (or contact-dependent signaling) is instead a type of cell signaling that requires close contact.

**Figure 6:**
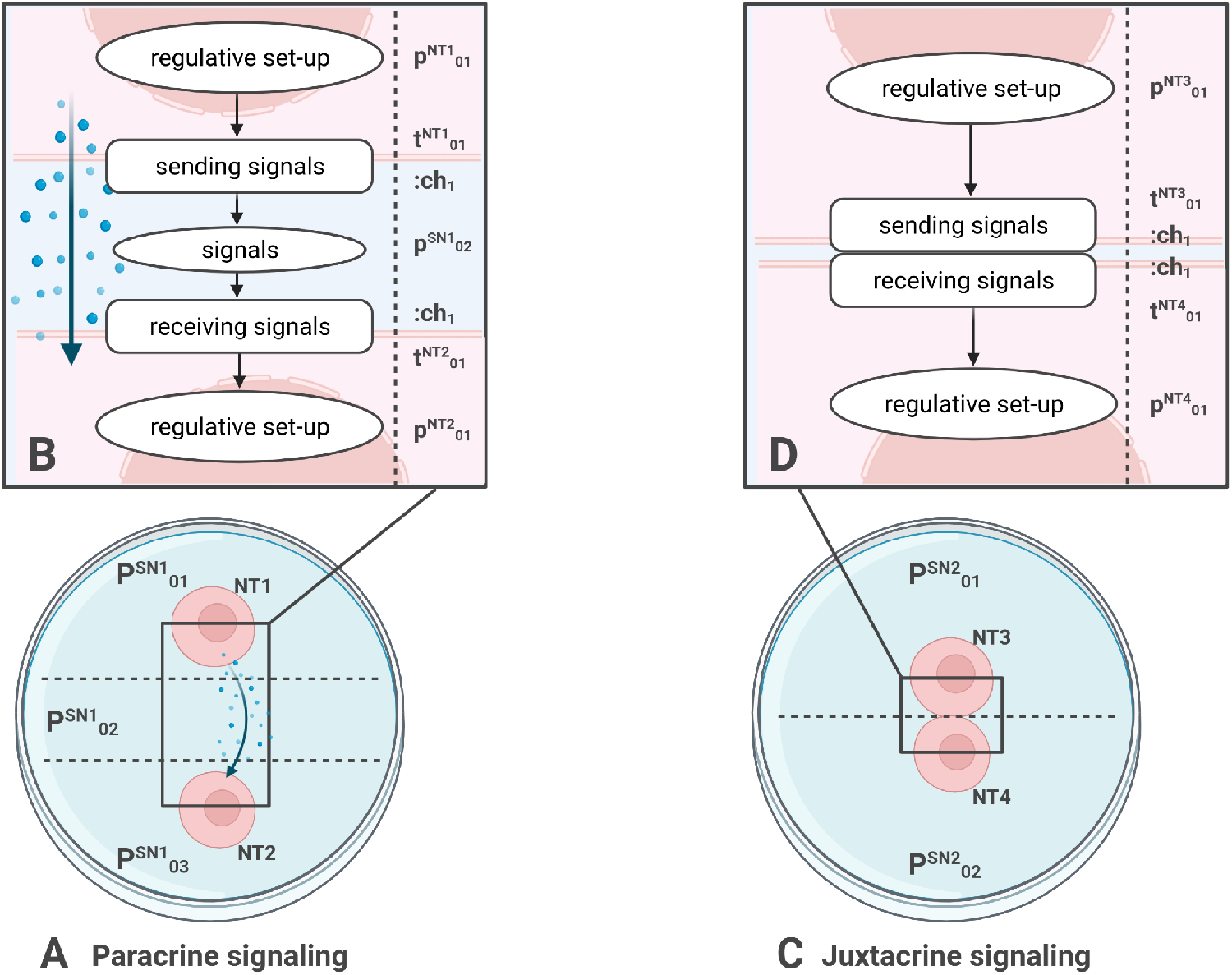
Paracrine and juxtacrine signaling. (A) Paracrine signaling between two cells exchanging a signal through the extracellular environment. (B) The regulative state of cell NT1 causes the cell to secrete a signal in the extracellular environment; cell NT2 receives the signal, which affects its regulative set-up. (C) Juxtacrine signaling between two cells exchanging signals through the intercellular space. (D) The regulative state of cell NT3 causes the cell to secrete a signal, which directly affects the regulative set-up of cell NT4.

The two communication mechanisms can be modeled according to the NWN reported in Figure 7. In paracrine signaling (Figure 7-A) a cell (NT1 in 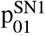) uses channel ch_1_ to send a signal represented by a black token in the extra-cellular environment 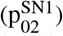. The extracellular environment here represented for simplicity as a single place can be also modeled with multiple places, based on the spatial organization of the cells. Another cell NT2 of a different type and in spatial proximity 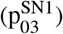 receives the signal using channel ch_2_. In juxtacrine signaling (Figure 7-B), two adjacent cells (NT3 and NT4 in 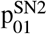 and 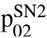, respectively) engage in direct communication through channels ch_1_ and ch_2_.

**Figure 7:**
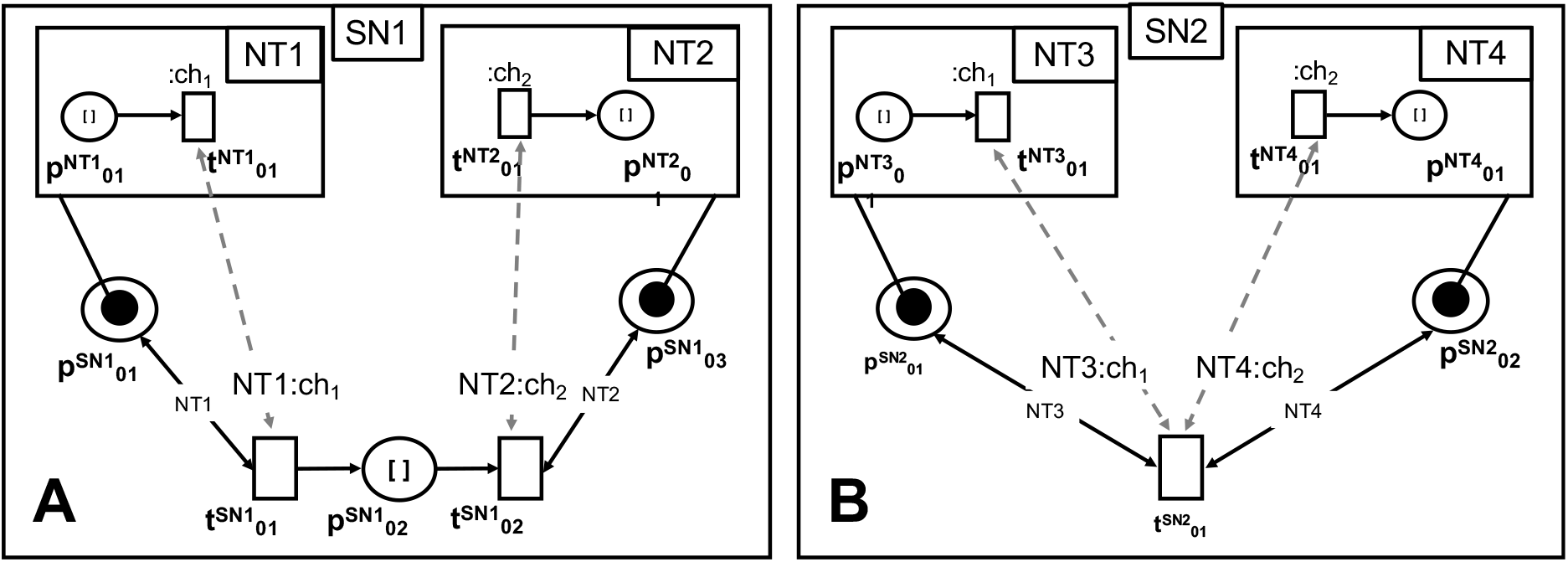
NWN representation of the two signaling mechanisms relevant for modeling ontogenetic processes: (A) In paracrine signaling, the cell NT1 uses channel ch_1_ to send a signal (black token) to the extracellular environment modeled as a place in the system-net 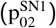. Cell NT2 that is in spatial proximity receives the signal using channel ch_2_. (B) In Juxtacrine signaling, the two adjacent cells (NT3 and NT4) engage in direct communication through channels ch_1_ and ch_2_.

These two network motifs are particularly relevant for modeling ontogenetic processes when combined with spatial information. During development, architectural complexity at the supra-cellular level emerges from local interactions between cells, which are often influenced by different distance ranges [52][51].

For example, sender cells engage in long-range interactions in paracrine communication, diffusing soluble signal molecules in the extracellular environment, targeting receiver cells. The signal starts with a higher concentration close to its source, and the concentration decreases with distance according to diffusion laws specific to the target molecule. Since the reaction to a signal is often dose-dependent, this mechanism translates into a distance-dependent effect over target cells.

In unilateral juxtacrine communication, sender cells communicate with receiver cells in their very proximity through their trans-membrane signal and receptor proteins, respectively. The cells involved in bilateral juxtacrine communication act through the same mechanisms, but each of them acts both as a sender and a receiver. Finally, in autocrine signaling, the cell sends out a signal intended to be self-received, acting both as a sender and as a receiver.

### 3.4. Dynamic hierarchy

The ability of recursively instantiating net-tokens within another net expresses the hierarchical and dynamic regulatory structures of biological systems (Figure 3.A). Let us consider the two-level model architecture of Figure 5:

- the system-net at the top-level describes the spatial organization and regulative landscape for two cells in two different process phases;
- the net-tokens existing as instances within the system-net represent cells living in such landscape.

One important aspect to highlight is that the hierarchy is not static but instead dynamically defined. At first, net-token instances are created in 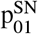 and 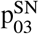, respectively. Instantiation per se does not carry any biological meaning, but the resulting marking reflects the system’s initial hierarchical organization. The system-net then evolves, and net-tokens can be created or destroyed, simulating the considered biological process, and modifying the system’s hierarchical architecture.

Moreover, NWN support the flexible specification of different abstraction layers (Figure 3-C). The same instance of a net-token can live into multiple higher-level nets, each one describing a different aspect of the considered process. This is an important characteristic that poses the basis for multilevel and multi-scale modeling.

Eventually, the cross-layer communication mechanisms presented in the previous sections provide a high degree of flexibility when deciding case by case which layer is in control of the evolution of a specific mechanism.

### 3.5. Process stages

Ontogenetic processes have sequential stages, with checkpoints involving regulatory states and sets of biological actors governing the passage from a stage to the next. Looking at Figure 5, most of the dynamics considered so far belong to the first of the two stages of the considered process. Transitions 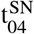 and 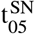 can transport net-tokens NT1 and NT2 to 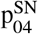 and 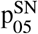, respectively. In this case, the transport represents the passage to a new regulatory setup. To occur, this passage needs some requirements at the net-token level to be satisfied. Both 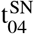 and 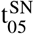 carry a down-link for the synchronous channel ch_2_. The up-link of this channel is in transition 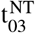 of the net-tokens. The channel creates an interplay between the two layers. To move a net token from 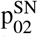 to 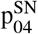 and therefore advance to the next process stage, it is not enough to have a net-token in a specific location of the system-net. The net-token must also satisfy an internal condition, i.e., one token must be present in 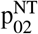. This condition represents the checkpoint for this developmental stage that models the gatekeeper mechanisms organizing ontogenetic processes into subsequent phases, i.e., some phenotypic traits need to be expressed by the cells for them to access the next stage. The ways each checkpoint evaluates net-tokens evolution and state can range from a simple read of a static value in the net to dynamic tracking to extract complex information about cell phenotype. The checkpoint may then include robust decision-making tools that implement classification routines that handle complex information and dynamically label complex net-token behaviors.

### 3.6. Dynamic regulatory landscape

The organization in different process stages also contributes to the definition of a dynamic regulatory landscape. If places 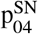 and 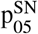 represent the counterpart of 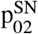 and 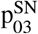 in a new developmental stage, the movement ofa cell (net-token) from one stage to another one changes the interaction schema that engages the cell. This modeling approach covers both intracellular (net-token) and supra-cellular (system-net) regulations, the latter intended as any regulation existing on top of intracellular regulation, including environmental and epigenetic context. Also, it allows expressing a priority scheme among such regulation layers and how that may change in different stages of the same process. This modeling approach allows representing a complex regulative landscape. Each point corresponds to a particular set of interactions that may take place in a specific process phase and context, and each cell can undergo multiple paths across it, passing through different process stages.

## 4. Results and discussion

In this section, we apply our modeling approach to a well-characterized developmental process: the Vulval Precursor Cells (VPC) specification in the development of *C. Elegans* larva. This aims to show the proposed modeling strategy at work, providing an overview of its capabilities. Differently from other models of the same process available in the literature (e.g., [9]), the proposed model includes different hierarchical levels, explicitly combines spatial information with cell differentiation and cell interaction, taking advantage of the proposed NWN modeling methodology.

### 4.1. Biological process

Vulval Precursor Cells (VPC) specification in *C. Elegans* is an ontogenetic process involving a small number of cells but still including all characteristics of more complex processes (Figure 8). Also, it is one of the most widely characterized processes of this kind at the intracellular level.

**Figure 8:**
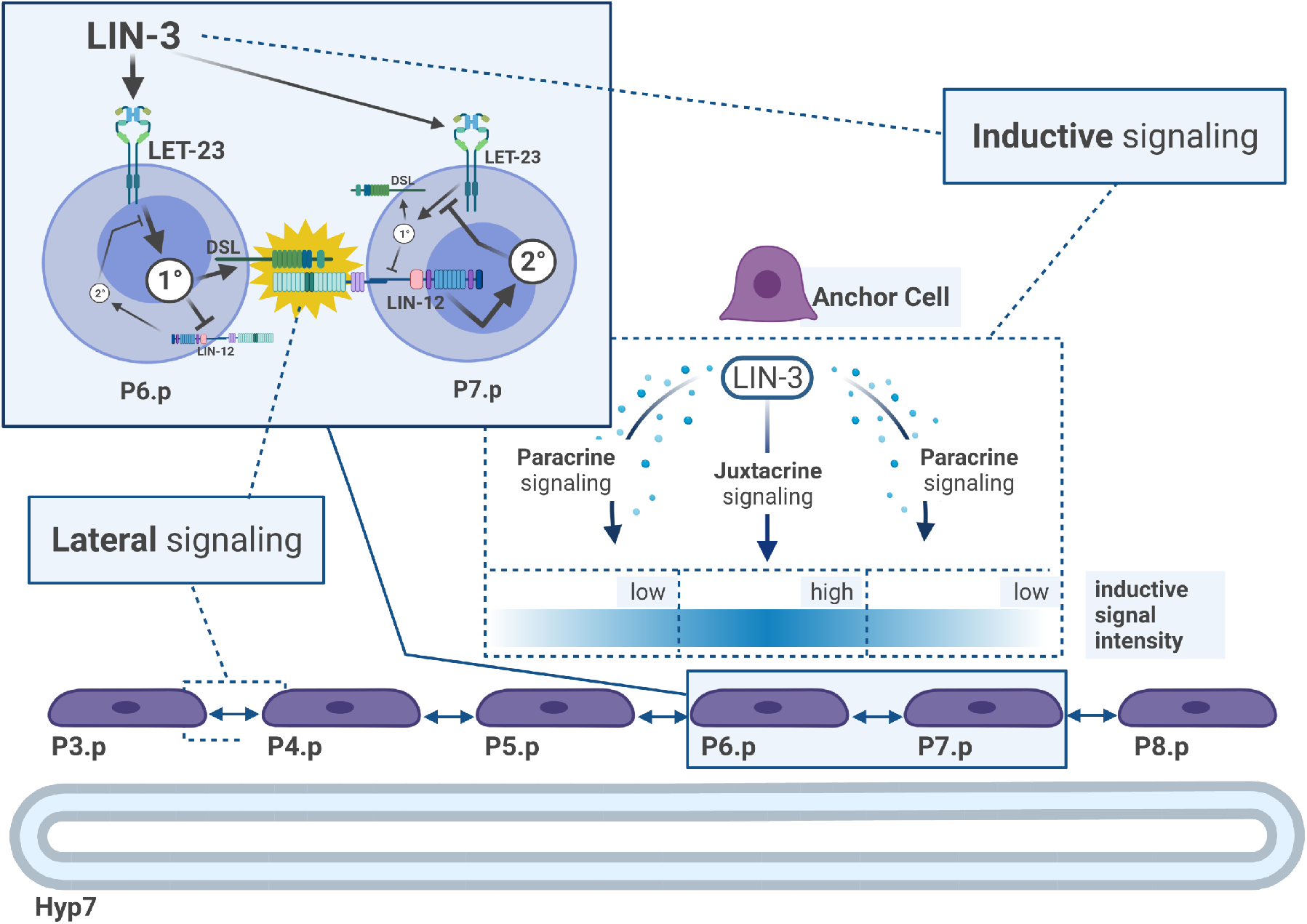
Diagram of signaling mechanisms involved in VPC pattern formation. During the L3 stage of C. Elegans larval development, inductive signaling from the Anchor Cell, belonging to the developing gonad, and lateral signaling among the VPCs (P3.p - P8.p) interact to make a precise pattern of cellular fates emerge. These include vulval lineages of two types, 1° and 2°, each of which generates distinct sets of progeny. The uninduced VPCs generate a 3° lineage, creating epidermal cells that fuse with the large syncytial epidermis hyp7. P6.p receives strong juxtacrine signaling via transmembrane LIN-3 signals from the AC. This strong signal induces the primary (1°) fate, suppressing the secondary (2°) fate and activating DSL lateral signaling to the two neighboring VPCs (P5.p and P7.p). P5.p and P7.p receive soluble, paracrine LIN-3 signaling from the AC, which is slower and weaker than juxtacrine signals and combines with the DSL signals from the neighboring P6.p to suppress the 1° and promote the 2° fates respectively. In the wild-type scenario, a precise spatial pattern of cellular fates emerges in the VPCs (3°-3°-2°-1°-2°-3°).

As extensively described in [69, 65], VPC specification occurs between the L3 and L4 stages of larval development in *C. Elegans*. At this stage, each of six multi-potent stem cells, the Pn.p cells, acquire one of three fates (1°, 2° or 3° fate), which guide the subsequent phases of organ development. Different actors contribute to fate decisions: the Anchor Cell (AC), residing in the adjacent developing uterus district, the underlying hypodermal syncytium, and the neighbor Pn.p cells.

More specifically, as in Figure 8, in the physiological case, the AC sends out a LIN-3 (EGF-like) signal reaching Pn.p cells with distance-dependent intensity: the closest cell, P6.p, receives juxtacrine signaling, its neighbors P5.p, and P7.p cells receive paracrine signaling. The signal does not reach P3.p, P4.p, and P8.p, the farthest cells. The hypodermal syncytium (hyp7) sends uniformly low-intensity paracrine LIN-3 signals to all Pn.p cells. The Pn.p cells can engage in mutual juxtacrine lateral signaling via trans-membrane DSL/LIN-12 (DSL/Notch-like) signaling. At the intracellular level, intense LIN-3 signaling induces the 1° fate in P6.p via the activation of a LET-23-mediated RAS/MAPK signaling pathway. This condition is marked by high concentrations of the active form of MPK-1, which activates strong DSL lateral signaling to the neighbors. This activation causes them to switch off the 1° fate traits induced by LIN-3 paracrine signals from the AC, activating 2° fate traits, corresponding to high concentrations of the active LIN-12 protein. P3.p, P4.p, and P8.p cells do not receive any LIN-3 other than that from hyp7, this causing them to undergo the 3° fate.

Different non-physiological cases for VPC specification exist due to various mechanisms, including genetic mutations.

### 4.2. The model

The NWN model of the VPC specification process comprises the following elements:

- a bi-dimensional Interactive Spatial Grid (ISG) (Figure 9);
- a Pn.p specific Differentiative Landscape (DL) (Figure 10) with complex checkpoint functionalities;
- three cell models, for the Anchor Cell, the Pn.p cells and the hypodermal syncytium (hyp7) (Figure 13, Figure 14 and Figure 15, respectively).

**Figure 9:**
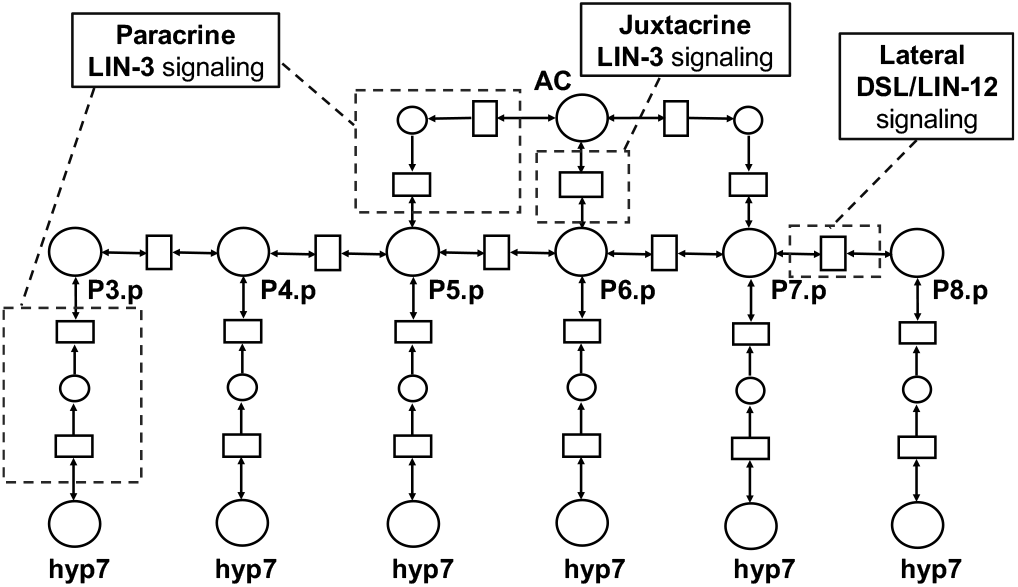
A schematic representation of the Interactive Spatial Grid (ISG) model for the VPC specification example, highlighting communication building blocks: *juxtacrine* LIN-3 signaling from the Anchor Cell (AC) to the P6.p cell; *paracrine* LIN-3 signaling from the Anchor Cell (AC) to the P5.p and P7.p cells; *neighbor* communication between pairs of Pn.p cells; *paracrine* LIN-3 signaling from the hypodermal syncytium (hyp7) to the Pn.p cells.

**Figure 10:**
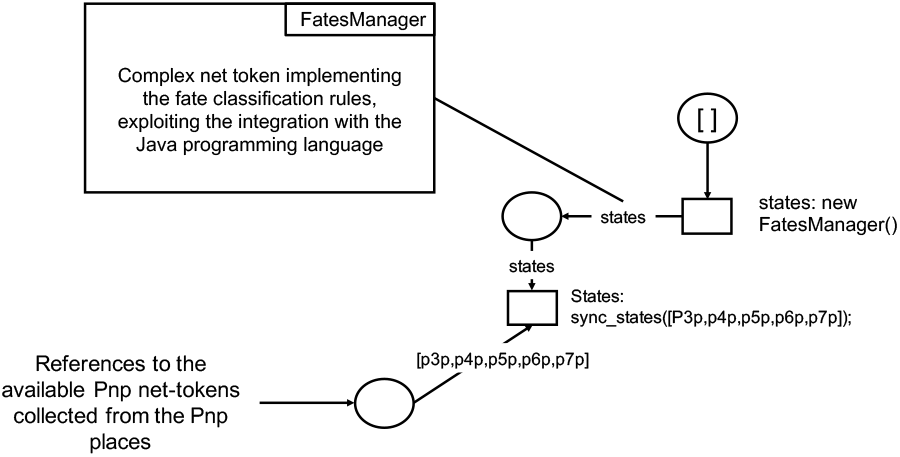
A schematic and simplified representation of the DL model for the VPC specification example. From the Pn.p places holding the cells, specific transitions create a complex token holding a reference to the Pn.p net-tokens. This array is passed to the FatesManager net-token using the sync_states channel. The Fates Manager is a special net-token that monitors the places modeling active MPK-1 and active LIN-12 of the Pn.p net-tokens and uses this information to predict the cell’s fate represented by assigning a color to the cells.

For readability, this paper resorts to simplified figures helping the reader understand the modeling approach. The complete model and instructions to reproduce the experiments are instead provided in a public GitHub repository at https://github.com/sysbio-polito/NWN_CElegans_VPC_model/.

Figure 11 helps to understand the hierarchical organization of the model’s elements. The system-net at the top-level combines ISG and DL functionalities. The nets that model the cells and implement the rules for classifying Pn.p cell fates (Fates Manager net) live and interact as net-tokens within this system-net.

**Figure 11:**
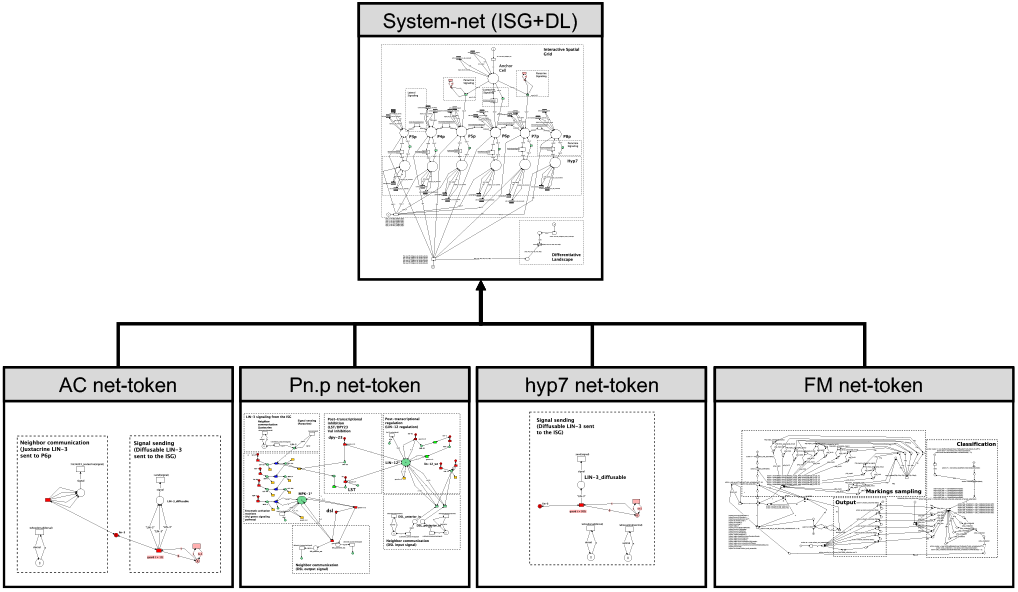
Nets hierarchy for the VPC specification model implementation in Renew. The system-net having ISG and DL functionalities is at the top level, and it hosts the net-tokens modeling different cells (AC, Pn.p, and hyp7), plus the Fates Manager (FM) modeling complex checkpoint rules.

Figure 12 reports a screenshot of the full system-net model implementation. The figure highlights the different portions this model using the dashed boxes, while the complete source file is available on GitHub.

**Figure 12:**
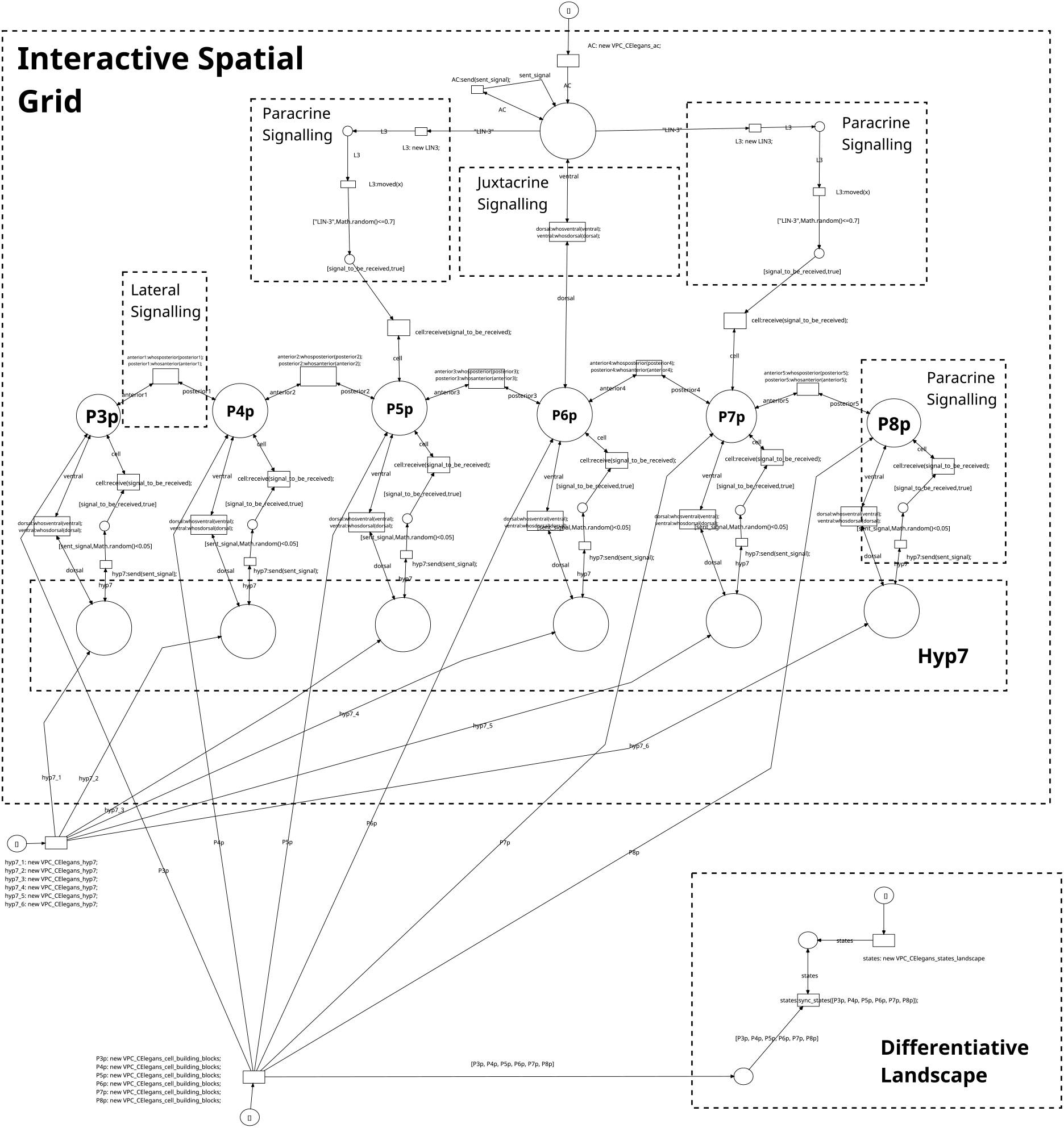
The Renew implementation of the system net for the VPC specification example, combining ISG and DL functionalities. This figure summarizes the complete architecture of the system-net to give an idea of the complexity of the model. A detailed view of this net is stored in the GitHub repository.

The top part of Figure 12 represents the ISG depicted in a more schematic way in Figure 9. It provides a two-dimensional representation of the VPC specification spatial environment and models the involved cells’ interactions. In this model, places represent the positions cells can occupy, and transitions model the potential interactions between cells in such positions (e.g., cell-cell communication mechanisms involved in the emergence of the morphogenetic pattern). Hyp7 sends LIN-3 paracrine signals uniformly to all Pn.p cells, modeling that the hypodermal syncytium lies under the array of Pn.p cells.

The DL model (Figure 10) represents instead the mechanism governing the Pn.p cells’ states along the developmental step from L3 to L4. Possible conditions are: Pn.p state, and Primary (1°), Secondary (2°), or Tertiary (3°) fates. Since this developmental stage does not imply a change in the system’s architecture, the different fates are modeled as colors of the net-tokens representing the Pn.p cells. Specific transitions from the Pn.p places holding the cells create an array of references to the Pn.p net-tokens. This array is passed to the Fates Manager net-token using the sync_state channel. The Fates Manager is a special net-token that monitors the places modeling active MPK-1 and active LIN-12 of the Pn.p net-tokens and uses this information to predict the cell’s fate. The bottom part of Figure 12 represents the actual implementation of this mechanism connected to the remaining parts of the system-net.

In the presented implementation, the system-net holds three main types of net-tokens.

The AC net-token describes the LIN-3 production and signaling in the Anchor Cell as reported in Figure 13. The AC model has two LIN-3 communication modes implemented using channels: one for *juxtacrine* signaling with P6.p (*neighbor communication*), and the other one for *paracrine* signaling with P5.p and P7.p (*signal sending*). Both signals rely on the LIN-3 gene for production (*transcription and translation*).

**Figure 13:**
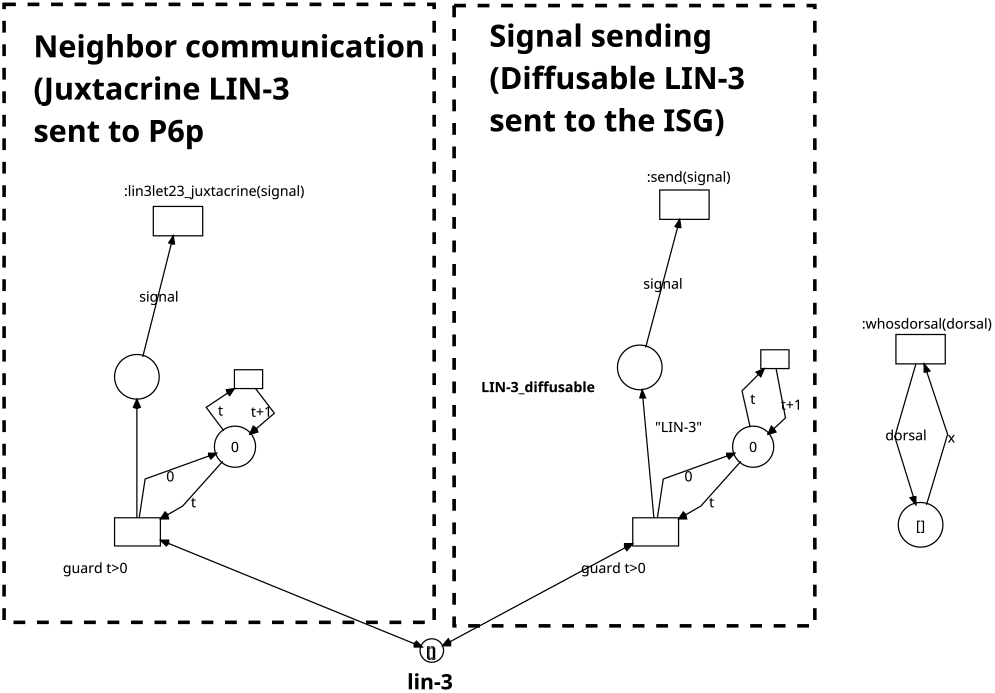
The Anchor Cell model. The Anchor Cell (AC) net-token models the two types of inductive signaling the AC sends to the Pn.p cells. A neighbor communication module (Appendix A, Figure 21) models juxtacrine signaling to P6.p, and a signal sending module (Appendix A, Figure 27) models paracrine signaling to P5.p and P7.p. Both signaling mechanisms leverage communication channels with the system-net (Paracrine and Juxtacrine signaling from the Anchor Cell place to the target Pn.p cells, Figure 12).

The Pn.p net-token describes the regulation of the Pn.p cells, all sharing the same architecture. This net-token is described using an adaptation of the Petri Nets model from [9]. As shown in Figure 14, the Pn.p cell model receives LIN-3 signals through channels that describe *paracrine* or *juxtacrine* signaling (*signal sensing* and *neighbor communication*). LIN-3 activates a cascade of enzymatic activation reactions (the MAPK signaling cascade, *enzymatic reactions*), resulting in the production of active MPK-1. This active protein causes the DSL signal to increase and affect neighbor cells. Pn.p Cells receive DSL signals from neighbors, and this activates LIN-12 (DSL/LIN-12 lateral signaling via *neighbor communication*). Increased active LIN-12 causes LST and DPY23 inhibitors to switch off the MAPK signaling cascade (*inhibitory post-transcriptional regulation*).

**Figure 14:**
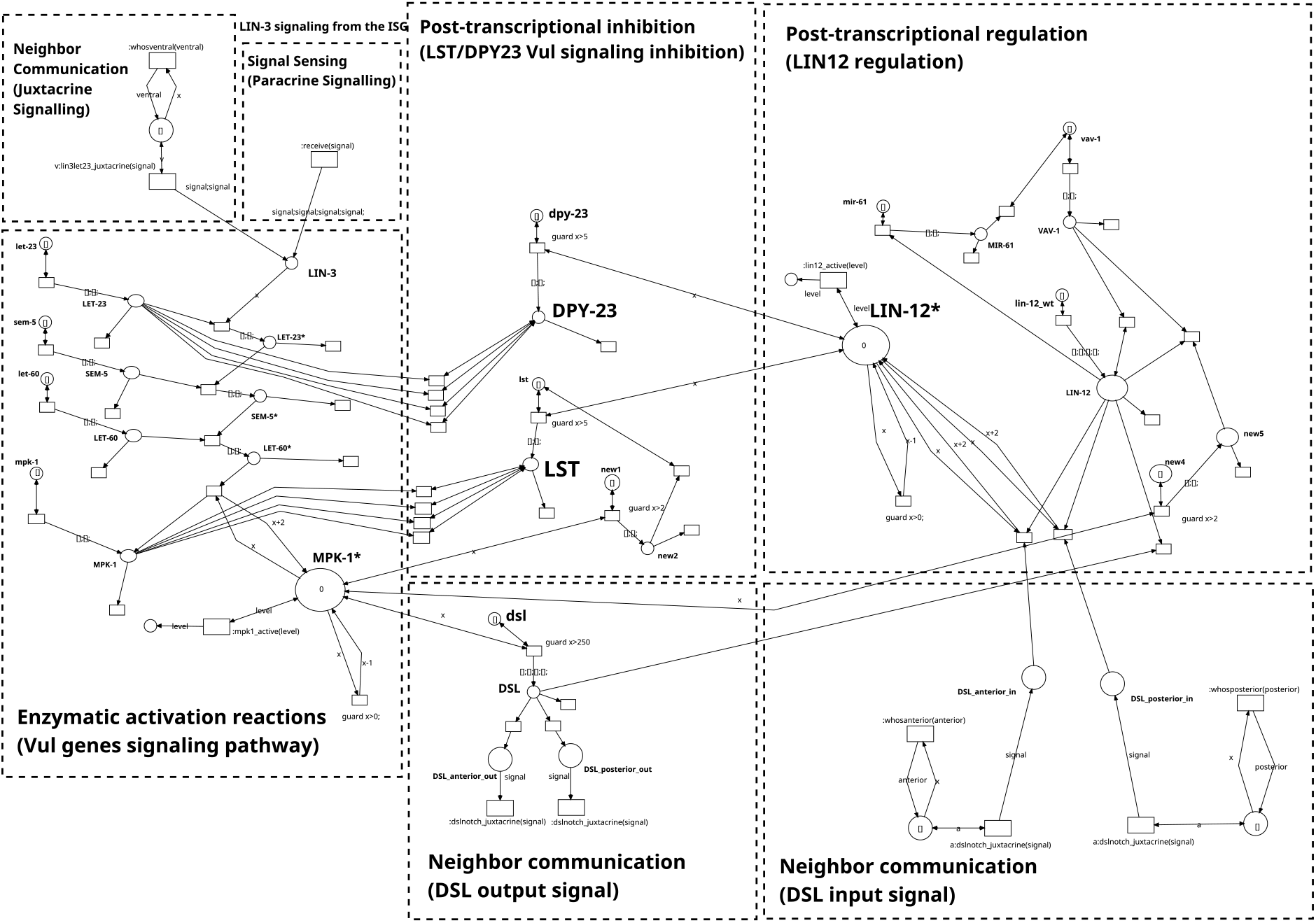
The Pn.p Cell model. The net-token for Pn.p cells models intracellular signaling cascades integrating inductive and lateral signals within each VPC. The AC communicates with Pn.p cells with either juxtacrine or paracrine inductive signaling that activates, with different intensities, the Vul genes signaling pathway. This pathway activates the 1° fate marker (i.e., active MPK-1) and the DSL output signal while inhibiting the 2° fate marker (i.e., active LIN-12) via DPY-23 and LST-mediated transcriptional inhibition. In addition, DSL-mediated lateral signaling activates the 2° fate marker (i.e., active LIN-12) in neighboring VPCs. LIN-12 is a Notch-like receptor for the Delta-like signal DSL. DSL-LIN-12 lateral signaling between neighbor Pn.p cells and AC juxtacrine signaling to P6.p cell are modeled with a neighbor communication module (see Appendix A, Figure 21). AC paracrine signaling is modeled combining a signal sending and a signal sensing module (Appendix A, Figure 27 and Figure 26).

**Figure 15:**
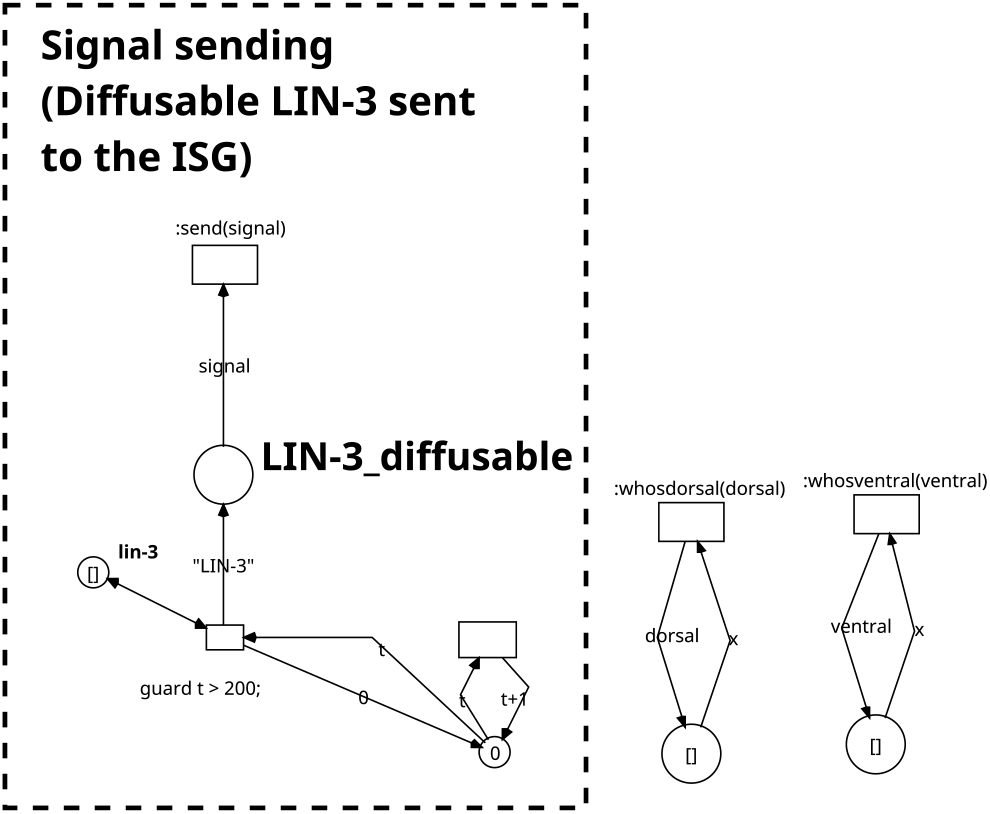
The hyp7 Cell model. This net-token models LIN-3 production by the hyp7 hypodermal syncytium with a signal sending module (Appendix A, Figure 26).

Eventually, the hyp7 net-token describes the hypodermal syncytium lying below the Pn.p cells. It provides uniform and low-intensity LIN-3 paracrine signals to all of them (*signal sending*), modeling the fact that LIN-15 can shut down LIN-3 transcription (*inhibitory transcriptional regulation*), as described in [9].

The Fates Manager (FM) net-token is a complex net able to assign one of the three possible fates (Primary, Secondary and Tertiary Fate) to the different Pn.p cells. The full network is available on GitHub. However, for the sake of readability, Figure 16 focuses on presenting the overall behavior of this network.

**Figure 16:**
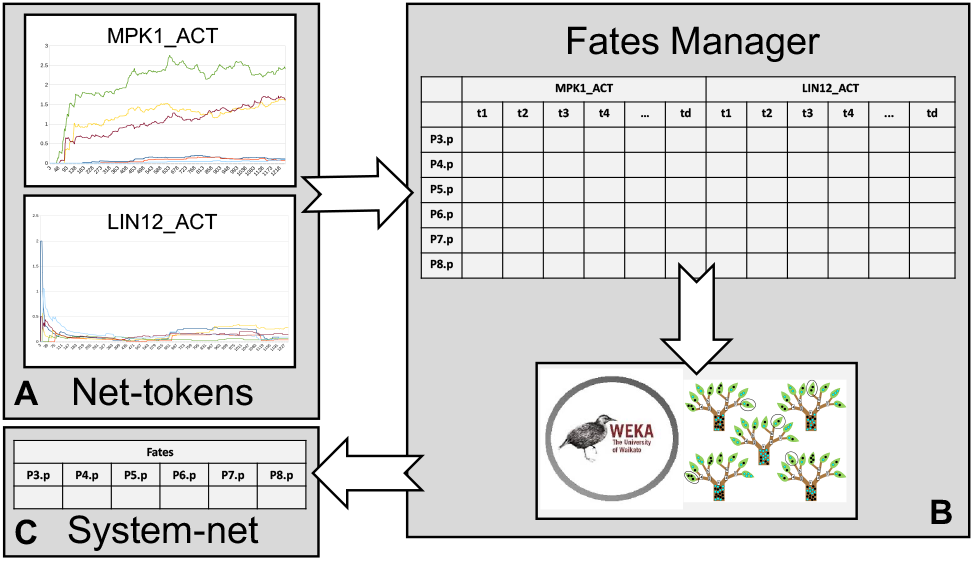
Overview of the cell fate classification process in the Fates Manager. The classification process relies on a Weka random forest classifier. The classifier receives as features information on the marking of the active MPK-1 (MPK-1_ACT and active LIN-12 (LIN-12_ACT) places of the Pn.p net-tokens and, based on it, predicts the cells’ fate.

The first task of the FM is sampling the marking evolution of the active MPK-1 and active LIN-12 places of all Pn.p cells. A channel transfers an array of Pn.p net-token object references to the FM that can then communicate with them to obtain their marking. The ability to track the dynamic evolution of places available in various instances of a net-token is a crucial characteristic of this modeling strategy. Apart from its usage in the presented model, these mechanisms can be used as a general and powerful inspection instrument to understand the considered process.

The FM stores the marking evolution of the active MPK-1 and active LIN-12 places of each cell into a set of circular buffers, creating a sliding window where the markings of the six Pn.p cells are stored during the simulation. This information builds a set of features for each specific cell submitted to a classifier able to determine the fate of the cell (Figure 16.B). Finally, the resulting fates are sent to the system-net (Figure 16.C) and used to assign the corresponding color to the Pn.p cells.

The classification process relies on a Weka random forest model [37]. The classification model is trained with a supervised mechanism. A training dataset is generated by collecting the marking of the active MPK-1 and active LIN-12 places in several simulations of the Pn.p net-tokens in a physiological condition. As will be detailed later in this section, simulations include stochastic behaviors that lead to different dynamics. Each set of features was manually labeled to a fate by comparing the evolution of the active MPK-1 and active LIN-12 markings with the expected values from biological knowledge. In particular, cells exhibiting high values of active MPK-1 were labeled with the Primary fate, cells having low active MPK-1 and high active LIN-12 with the Secondary fate, and cells having low levels of both markers with the Tertiary fate.

### 4.3. Model tuning and simulations

The proposed implementation of the Pn.p, AC, and hyp7 net-tokens follows the semi-quantitative paradigm proposed in [9]. The tuning proceeded manually by trial-and-error until the marking evolution of markers (i.e., active MPK-1 and active LIN-12) recapitulated physiology for all cell models.

Renew supports different high-level Petri Nets formalisms and their simulation. Experiments use the Java Net Compiler [11], which supports stochastic simulation of NWN models. The simulation engine considers all enabled transitions at each cycle and then randomly selects a subset of them for activation. This mechanism makes the transitions activation order non-deterministic, mimicking the stochasticity of the biological processes. In other words, this means that starting from the same initial conditions, simulations of the same model may generate different outcomes.

Moreover, the model structure introduces delays for some mechanisms to provide better expressivity of graded signaling intensity. For example, expressing the delayed onset of *paracrine* signals from the AC compared to the *juxtacrine* ones.

This modeling approach expresses temporal dynamics, which are relevant for modeling ontogenesis since the timing of functional activation often has a regulatory meaning in that context.

Experiments involved two simulation campaigns. The first simulation campaign aimed at collecting features to train the FM classifier. In this campaign, the Weka classifier was detached from the model. The FM only collected the features from the marking of the Pn.p cells and stored them in a file (see training_set.arff file on GitHub). Collected data were then labeled and used to train the model. Data from simulations were enriched with synthetic data to cover corner cases difficult to obtain from raw simulations. Globally, the training set contains 100 samples for each of the three possible fates.

The second campaign shows the model at work. Different initial markings represent different experimental conditions in the model, chosen by the scheme proposed in [9]. In particular, this paper considers the following conditions:

1. *wt*: the wild-type physiological condition;
2. *lin12_ko*: knock-out of the lin-12 gene;
3. *lstdpy_ko*: knock-out of the lst (lip-1, lst-1, lst-2, lst-3 and lst-4) and dpy23 set of genes ;
4. *vul_ko*: knock-out of the Vul set of genes (let-23, sem-5, let-60 and mpk-1).

Each condition corresponds to an expected VPC differentiation pattern as follows:

1. *wt*: P3.p: 3° fate, P4.p: 3° fate, P5.p: 2° fate, P6.p: 1° fate, P7.p: 2° fate, P8.p: 3° fate;
2. *lin12_ko*: P3.p: 3° fate, P4.p: 3° fate, P5.p: 1° fate, P6.p: 1° fate, P7.p: 1° fate, P8.p: 3° fate;
3. *lstdpy_ko*: P3.p: 3° fate, P4.p: 3° fate, P5.p: 1° fate, P6.p: 1° fate, P7.p: 1° fate, P8.p: 3° fate;
4. *vul_ko*: P3.p: 3° fate, P4.p: 3° fate, P5.p: 3° fate, P6.p: 3° fate, P7.p: 3° fate, P8.p: 3° fate;.

For each experimental condition, 100 simulations were collected following recommendations in [60]. Simulations provide both the temporal evolution of the markers of interest from cell models and the six labels resulting from Pn.p cells classification.

As mentioned before, validation and tuning of the model were performed using simulations. During simulation, the temporal evolution of the marking of each place of the different networks composing the model was constantly recorded. The marking of a place can be seen as the intensity of the biological signal associated to the place at a given point in time. The average marking over a complete simulation represents the average signal intensity. This metric, after min-max normalization, is used here to analyze and validate the behavior of the model by looking at a sample simulation from the *wt* condition. Figure 17 focuses on the dynamics of the P5.p, P6.p, and P7.p cells. As expected from the knowledge of the biological system, the P6.p cell receives a strong LIN-3 juxtacrine signal from the AC that corresponds to significant active LET-23 production (0.25 normalized average intensity), which induces a strong production of active MPK-1 (0.5 normalized average intensity), inducing the Primary fate. At the same time, the P6.p cell does not receive significant DSL lateral signaling from the neighbors, resulting in a negligible production of active LIN-12. Both the P5.p and P7.p cells receive weaker paracrine LIN-3 signal from the anchor cell, which requires some time to be delivered. This corresponds to a weaker production of active LET-23 (less than 0.1 normalized average intensity) that starts later than the P6.p cell. Moreover, these cells receive strong DSL lateral signaling (about 0.25 normalized average intensity) induced by high active MPK-1 production in the P6.p cell. This results in a strong production of active LIN-12 (about 0.4 normalized average intensity) that inhibits the production of active MPK-1 (about 0.2 normalized average intensity), a condition that correctly corresponds to the Secondary fate expected to be induced in these cells.

**Figure 17:**
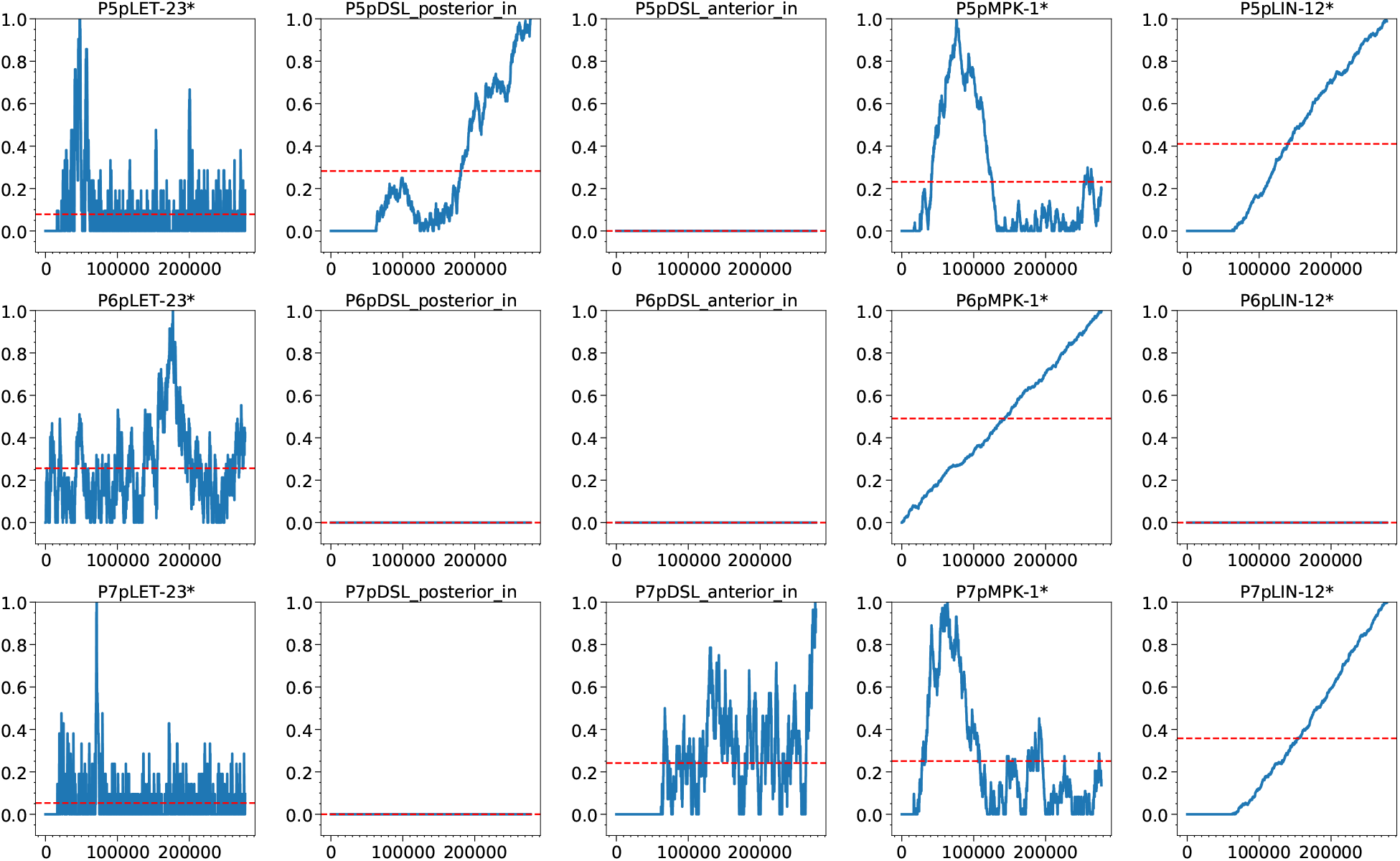
P5.p, P6.p, and P7.p dynamic behavior. The dynamic marking evolution of relevant places along with the simulation in the Pn.p models correctly recapitulates physiology. P6.p (central row) receives strong LIN-3 signaling from the AC, resulting in strong MPK-1 activation and DSL-mediated lateral signaling to neighbors. P5.p and P7.p (top and bottom rows, respectively) receive weaker, delayed LIN-3 signal, inducing weak LET-23 activation, leading to moderate active MPK-1 levels, which are inhibited by the high LIN-12 activation induced by the DSL lateral signaling from the P6.p cell, inducing the Secondary fate.

Figure 18 extends the analysis to all Pn.p cells. As expected cells P3.p, P4.p and P8.p (first, second and last rows, respectively) only receive weak LIN-3 signal from the hypoderm and may receive very weak lateral signaling from neighbors. This condition is not enough to trigger a significant active MPK-1 nor active LIN-12 production. Therefore, in these cells, both markers levels are negligible (around 0.0 normalized average intensity), which indicates the Tertiary fate.

**Figure 18:**
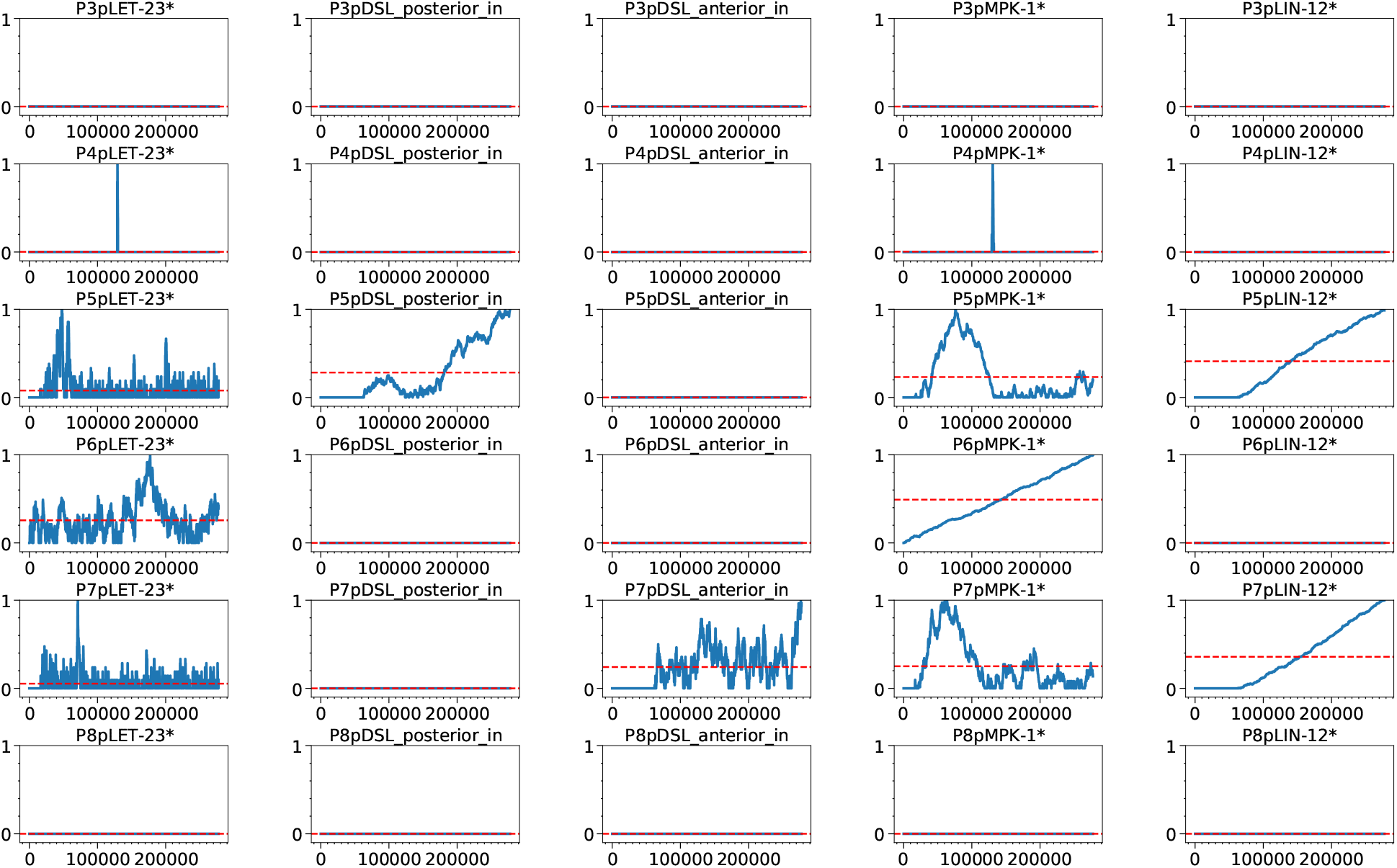
Pn.p cells dynamic behavior. The dynamic marking evolution of relevant places in the Pn.p models along with the simulation correctly recapitulates physiology. P6.p (fourth row) receives strong LIN-3 signaling from the AC, resulting in strong MPK-1 activation and DSL-mediated lateral signaling to neighbors, simulating the onset of the Primary fate. P5.p and P7.p (third and fifth rows, respectively) receive weaker, delayed LIN-3 signal, inducing weak LET-23 activation, leading to moderate active MPK-1 levels, which are inhibited by the high LIN-12 activation induced by the DSL lateral signaling from the P6.p cell, inducing the Secondary fate. P3.p, P4.p, and P8.p (first, second, and last row, respectively) do not receive any LIN-3 inductive signaling from the AC, nor do they receive significant DSL lateral signaling from neighbors, resulting in negligible levels of both markers active MPK-1 and active LIN-12, which correctly recapitulates their Tertiary fate.

While Figure 17 and Figure 18 show the dynamic behavior of the system analyzing a single simulation in the wildtype physiological condition, Figure 19 shows the results of several simulations on all considered conditions, represented as the normalized average values of the active MPK-1 and active LIN-12 markers for simulated Pn.p cells. The figure shows the ability of the model to recreate, for each considered condition, the expected differentiation pattern influenced by the corresponding level of the active MPK-1 and active LIN-12 markers.

**Figure 19:**
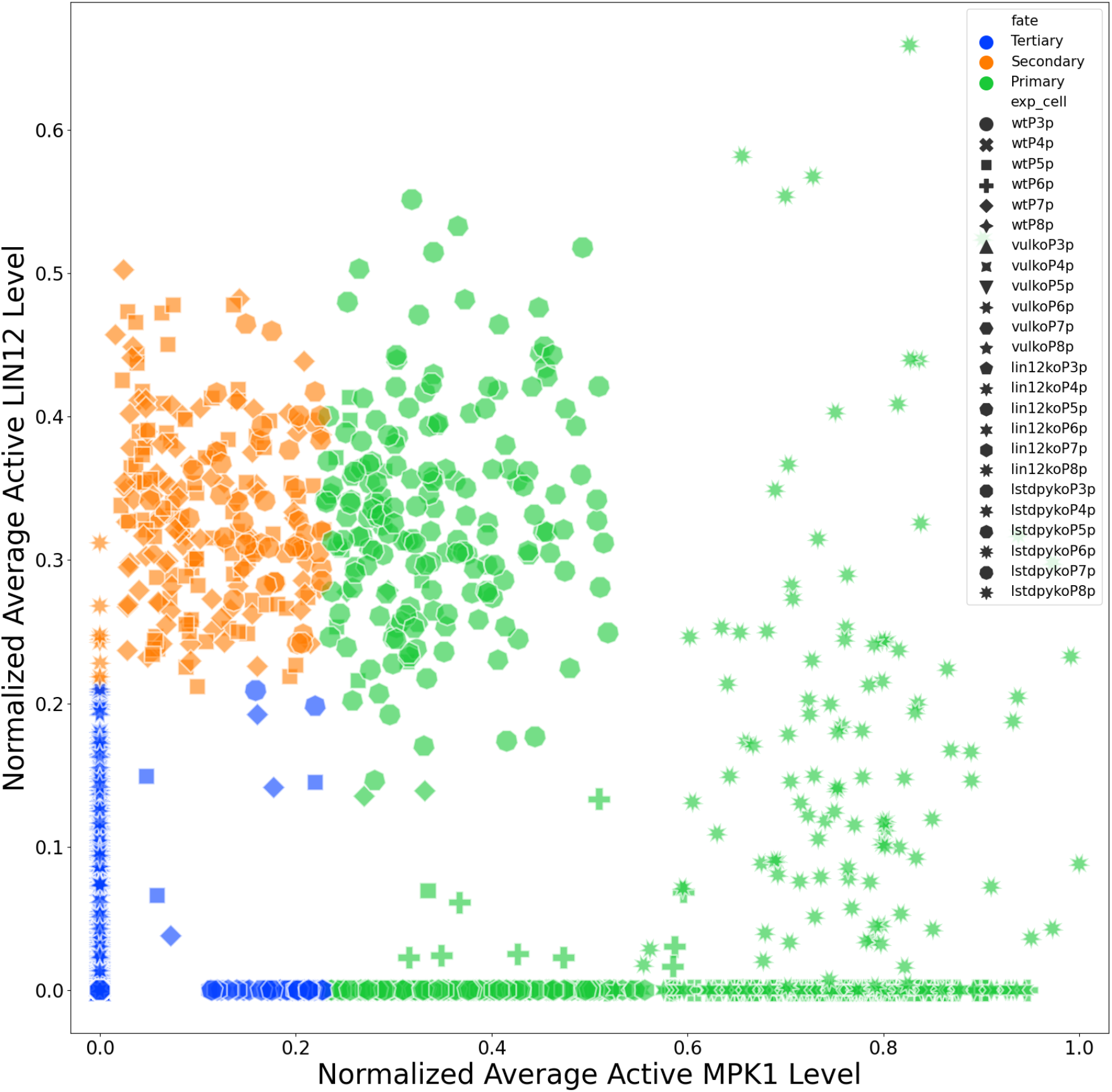
Simulations results. This plot illustrates that simulations from different experimental conditions (*wt, vul_ko, lin12_ko* and *lstdpy_ko*) correctly recapitulates the expected biological behavior in terms of markers levels and assigned fates for the Pn.p cells.

Of course, misclassifications are possible for simulations generating signals close to the classifier’s boundary. This is possible due to the stochasticity of the simulation. To quantify the accuracy of the model, Table 1 summarizes, for each cell, the ability of the model to predict the correct fate of the different cells in the considered conditions. These values refer to 100 simulations per condition. It is important to remark that the model was trained and tuned on the *wt* case and then used to evaluate new unknown conditions. This is a simple example to show how this modeling strategy can be exploited to inspect different conditions, therefore, performing *in silico* knowledge exploration.

**Table 1.** Results of the pattern formation simulations. For each experimental condition, the Pn.p fates pattern observed is reported, together with the accuracy of the prediction for each cell generated in the simulation outcomes. Reported results refer to 100 simulation runs.

| experiment | Pn.p pattern | accuracies |
| --- | --- | --- |
| wt | 3 3 2 1 2 3 | 100% 100% 87% 100% 93% 100% |
| lin12_ko | 3 3 1 1 1 3 | 100% 97% 74% 100% 83% 100% |
| lst_lf | 3 3 1 1 1 3 | 100% 100% 79% 100% 83% 100% |
| vul_ko | 3 3 3 3 3 3 | 100% 100% 100% 100% 100% 100% |

## 5. Conclusions and future perspectives

This paper presented an NWN-based modeling approach responding to modeling requirements posed by ontogenesis applied to the case study of a VPC specification in C. Elegans. Considering model performance in terms of predictive power, compared to other state-of-the-art approaches [9, 18, 38, 25], our modeling framework has similar or better predictive performance. Furthermore, the proposed model has the advantage of supporting the integration and composition of heterogeneous biological information and models. It combines the advantages of multi-level hybrid models [4] with formalism uniformity, which facilitates model analysis, and knowledge representation and exchange.

In the future, performance needs to be tested over a more significant number of mutations for the VPC specification case, starting from those tested in [8] and performing predictions over new experimental conditions to be subsequently verified experimentally in order to demonstrate the actual predictive power of the model.

At the moment, NWN models have been mainly used through simulation since the formal analysis of such complex graphs is not trivial due to the state-space explosion problem [30]. In particular, model checking is a promising method to verify systems that are modeled as state transition graphs [14]. Moreover, besides the verification capabilities, formal methods can support additional analysis able to infer system level properties of the system (e,g, invariants, stead states, etc.) that could provide interesting insights on the studied phenomenon.

Low-level PN are supported by a large literature providing methods like partial order reduction [72], symmetries [66], the sweep-line method [13] or alternative ways to represent the state space [20, 55] and cope with large model sizes. Tools such as Maria [47], LoLA [75], GreatSPN [1], Maude [15] are examples of instruments providing verification options for traditional PN variants. For high-level Petri nets some interesting model checking tools also exist [16, 21, 32]. However, the problem is still open when considering the peculiarities of the NWN formalism. Venero et al. [73] provide a method that can be applied to multilevel and recursive nets. Recently, Willrodt et al. [74] presented Modular Model Checker (MoMoC), a model checking tool designed to work with NWN and integrated with Renew. While this is a promising starting point, these tools can handle only very simple nets, and significant work is still required to handle complex models such as the ones presented in this paper.

Our models combine hypothesis-driven and data-driven approaches, but a necessary improvement is to make parameter identification rely on experimental data only, possibly generating them *ad hoc*.

Computational capabilities could limit the model complexity: if computational complexity is too high, analyses and simulations take too long to complete [54]. However, it is possible to face this problem both at the model level, with complexity reduction [2], and at the hardware level, with state-of-the-art architectures for parallel and distributed computing to speed up computational times [26, 28].

Finally, significant effort is required to improve the modeling strategy usability from life scientists with limited experience in model development and computer engineering. An effort in this direction is already in place with the development of the Biological System Description Language (BiSDL), a domain-specific high-level, modular programming language naturally accessible by purely biological semantics and automatically generating simulation-ready NWN models [49].

In this way, we intend to improve the capability of modeling biological complexity and make the resulting tools available to a large and diverse user base to fulfill the systems biology scientific community’s needs [58].

---

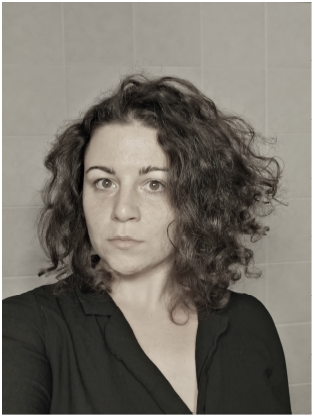

Roberta Bardini is a post-doc researcher at Department of Control and Computer Engineering of Politecnico di Torino. Her research focuses on computational approaches to understand and represent complex biological systems.

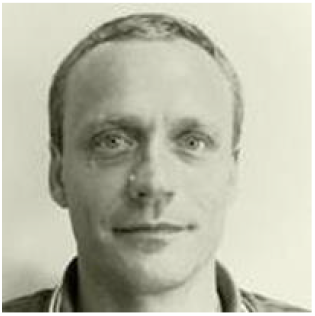

Alfredo Benso received the MS degree in computer engineering and the PhD degree in information technologies, both from Politecnico di Torino, Italy, where he is working as a tenured associate professor of computer engineering. His current research interests include bioinformatics, and system biology. He is also actively involved in the Computer Society, where he has been a leading volunteer for several projects. He is a Computer Society Golden Core Member, and a senior member of the IEEE.

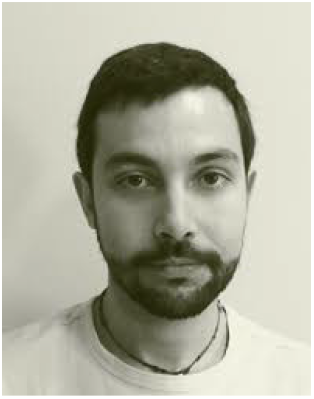

Gianfranco Politano is an Assistant Professor in the department of Control and Computer Engineering at Politecnico di Torino (Italy). He holds a Ph.D. and an M.S. equivalent in Computer Engineering and Information Technology from the Politecnico di Torino. Politano’s research contributions include complex biological networks analysis, data fusion and statistical data analysis.

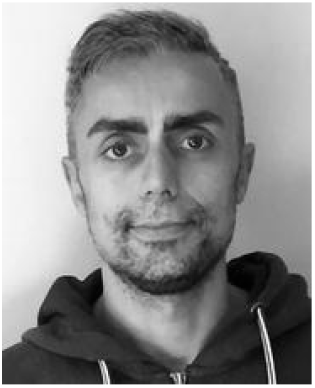

Stefano Di Carlo is an associate professor in the department of Control and Computer Engineering at Politecnico di Torino (Italy) since 2014. He holds a Ph.D. (2003) and an M.S. equivalent (1999) in Computer Engineering and Information Technology from the Politecnico di Torino in Italy. Di Carlo’s research contributions include biological network analysis and simulation, machine learning, image processing and evolutionary algorithms, as well as Reliability, Memory Testing and BIST.

## A. Appendix 1

This appendix reports a library of biological functional modules and their respective NWN model that can be used to model complex ontogenetic processes.

### A.1. Interactive spatial grid

This section reports a set of cell movement and cell to cell communication mechanisms mediated by the relative spatial positions modeled in the ISG.

**Figure 20:**
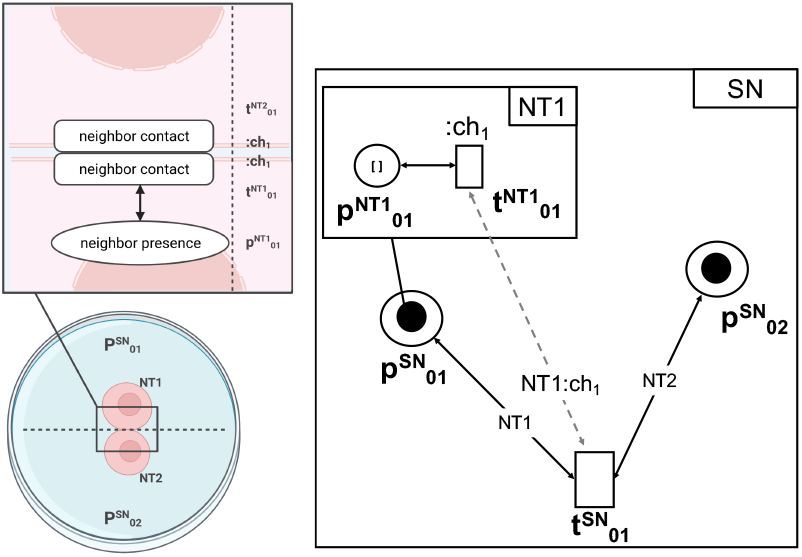
Neighbor detection. It models the capability of a cell to detect the presence of a neighbor cell in direct physical contact. This module allows a net-token occupying a place in the ISG (NT1 in 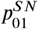) to detect a neighboring net-token instance in the adjacent place 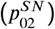 via a communication channel :*ch*_1_, linking transitions 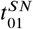 in the system-net (SN) and 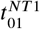 in NT1.

**Figure 21:**
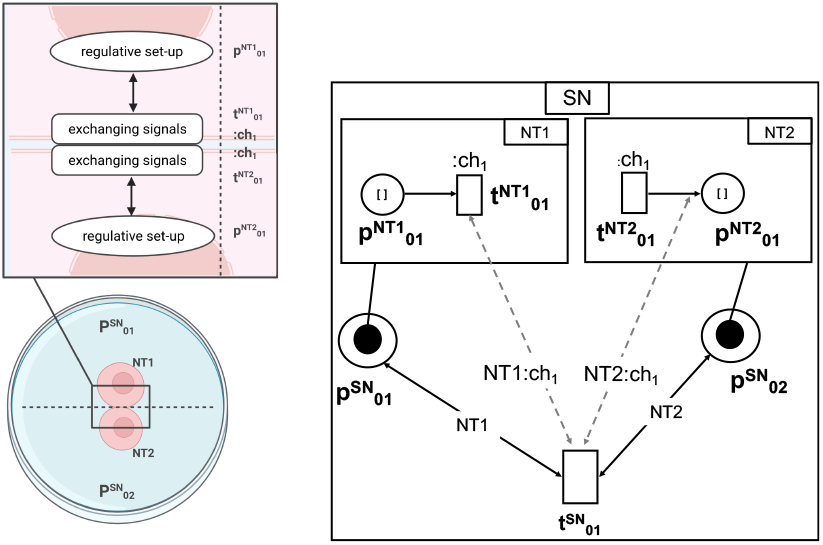
Neighbor communication. It models the communication between neighboring cells in direct physical contact. This module allows a net-token occupying a place in the ISG (NT1 in 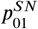) to connect to the net-token instance living in the neighboring place (NT2 in 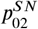) by exchanging signals via a communication channel :*ch*_1_, linking transitions 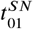 in the system-net (SN), 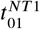 in NT1 and 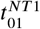 in NT2.

**Figure 22:**
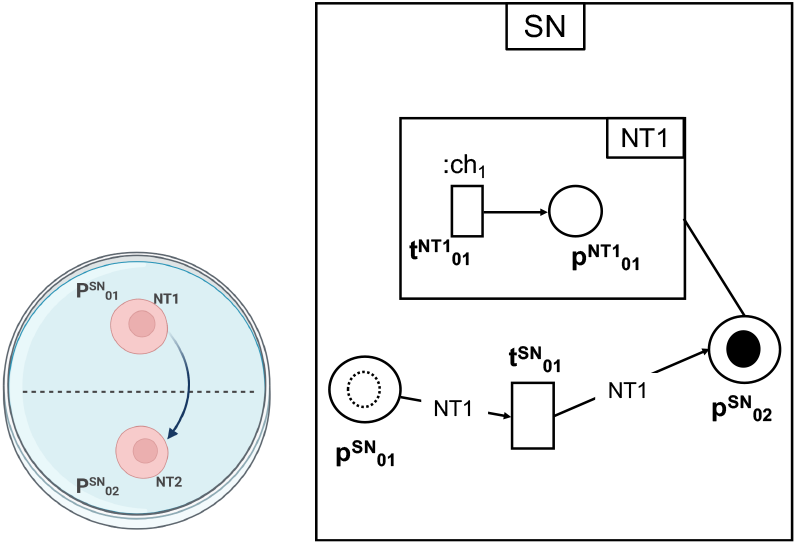
Cell movement. It models the mobility of a cell from a one position to another one in the physical environment. This module describes the movement of a net-token (NT1) from a place in the ISG 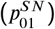 to another one 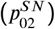 via a transition 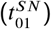 in the system-net (SN).

**Figure 23:**
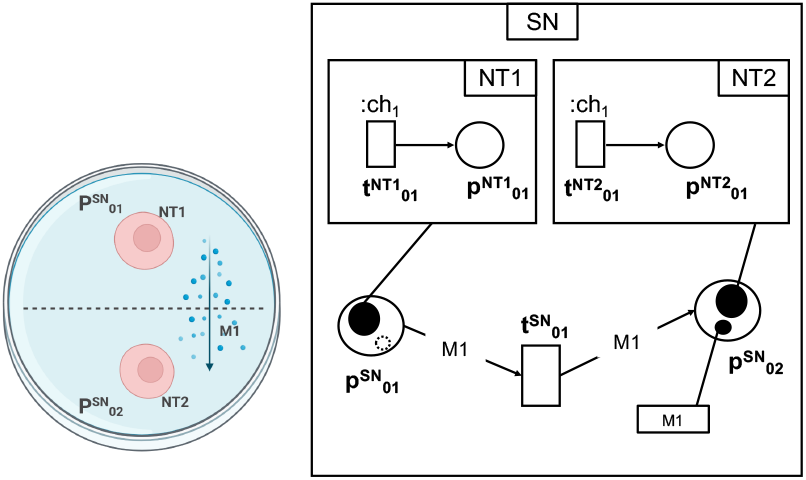
Molecular flow. It models the mobility of a molecule from a one position to another one in the physical environment. This module models the movement of a colored token (M1) from a place in the ISG (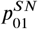, holding the net-token NT1) to another one (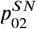, holding the net-token NT2) via a transition 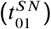 in the system-net (SN).

**Figure 24:**
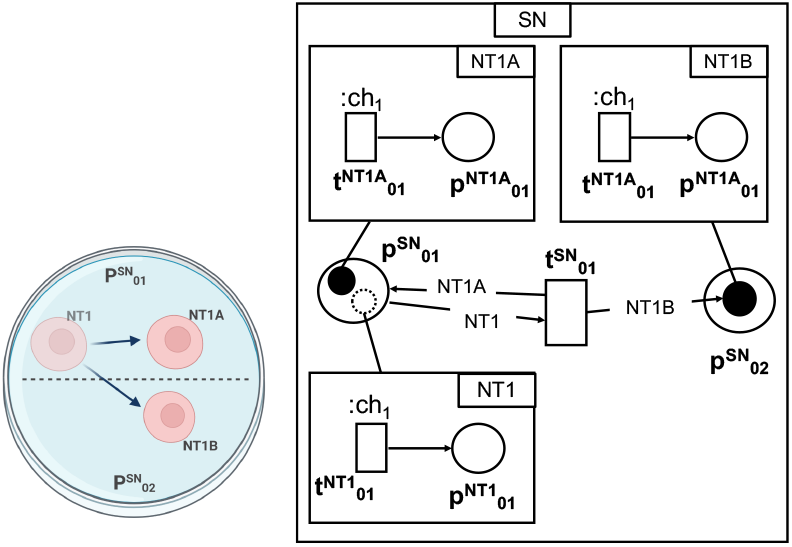
Mitosis. It models a cell undergoing mitosis and generating two daughter cells. This module includes the mother cell (NT1) in a position in the ISG 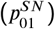 generating two daughter cells via a transition 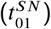 in the system-net (SN), one in its starting position 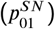, and another one in close spatial proximity 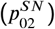.

**Figure 25:**
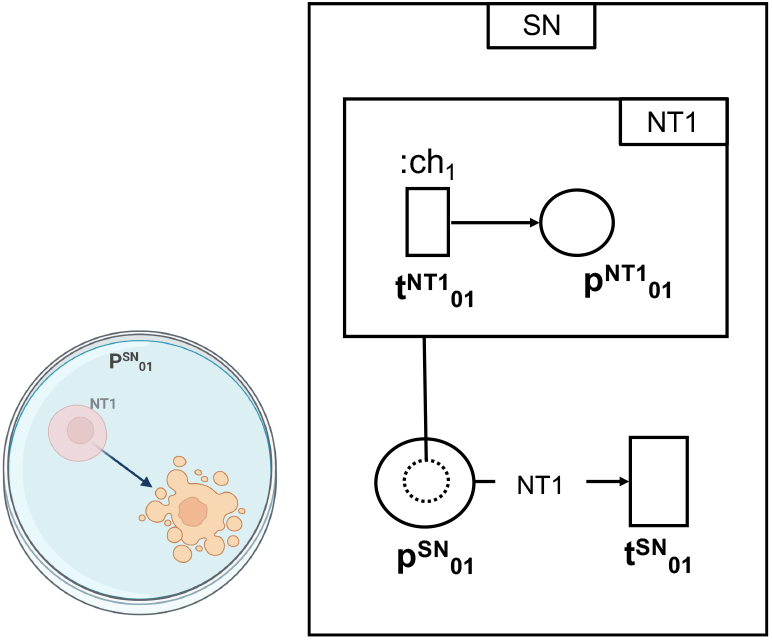
Apoptosis. It models a cell undergoing apoptosis that leaves the position it occupied in the environment empty. This module includes the apoptotic cell (NT1) in a position in the ISG 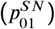 being removed from the ISG via a transition 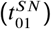 in the system-net (SN).

**Figure 26:**
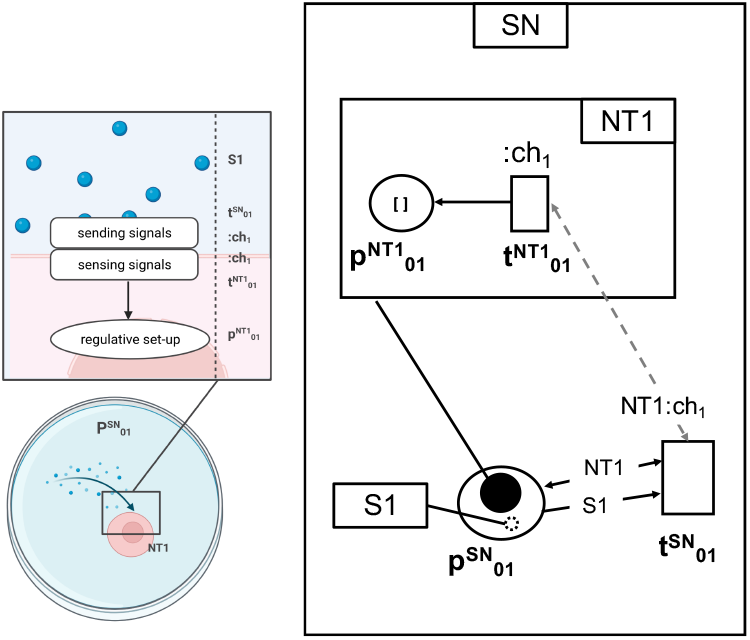
Signal sensing. It models a cell sensing a signal from the extracellular environment. In this module, the net-token occupying a place in the ISG (NT1) senses the signal modeled with the colored token S1 in the same place 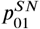 it occupies via a communication channel :*ch*_1_, linking transitions 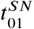 in the system-net (SN) and 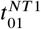 in NT1.

**Figure 27:**
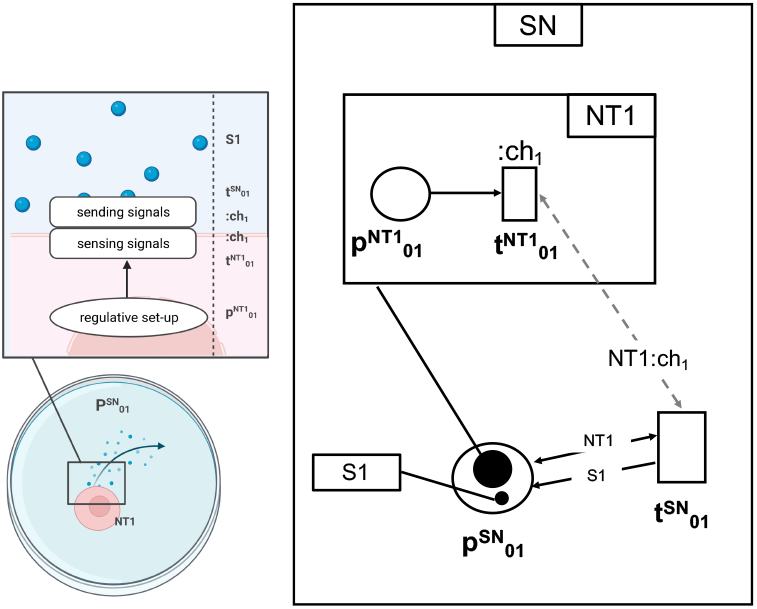
Signal sending. it models a cell sending a signal to the extracellular environment. In this module the net-token NT1 sends the signal modeled with the colored token S1 in the same place it occupies in the ISG 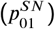 via a communication channel :*ch*_1_, linking transitions 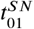 in the system-net (SN) and 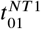 in NT1.

### A.2. Differentiative landscape

This section reports a building block to model a single differentiative step in a Differentiative Landscape (DL) model.

**Figure 28:**
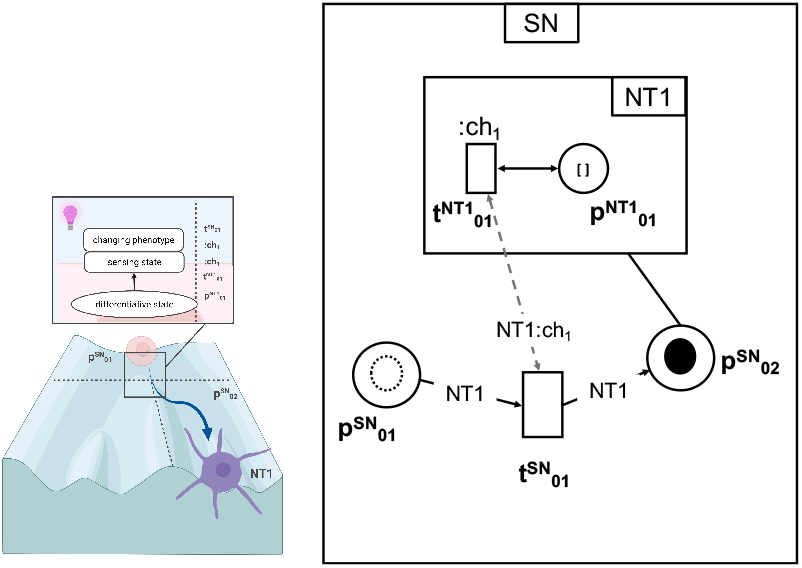
Differentiative step. It models a cell changing its phenotype state. This passage is regulated by a checkpoint evaluating the cell state. This module includes the net-token occupying a place in the DL (NT1 in 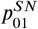). If NT1 internal state activates the checkpoint in 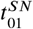 via a communication channel :*ch*_1_ (linking 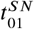 in the system-net and 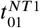 in NT1), the DL transition moves NT1 to 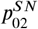, modeling the new phenotypic state.

### A.3. Cells

This section reports a set of sample building blocks to model simple intracellular mechanisms in the Cell models.

**Figure 29:**
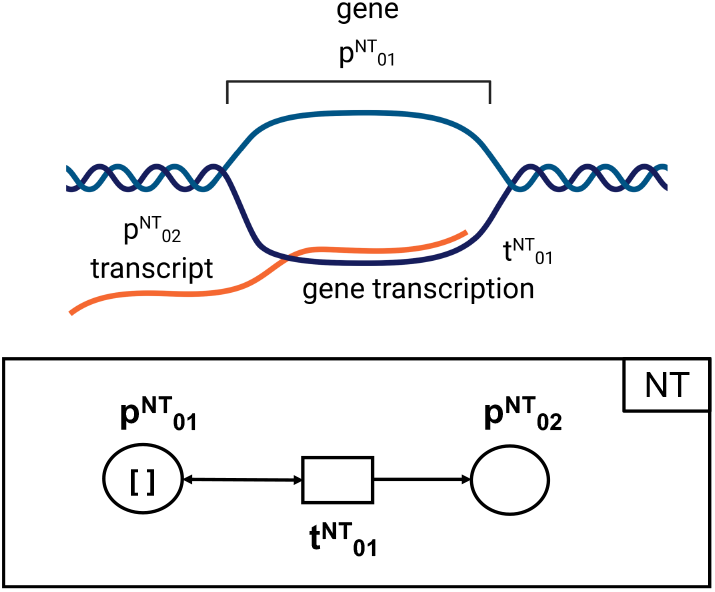
Transcription. It models the transcription of a gene (or a region of the genome) into coding or non-coding RNA transcripts. This module includes a net-token (NT) place for the genomic information to be transcribed 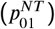 and one for the transcription products 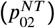, while 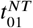 models the transcription process.

**Figure 30:**
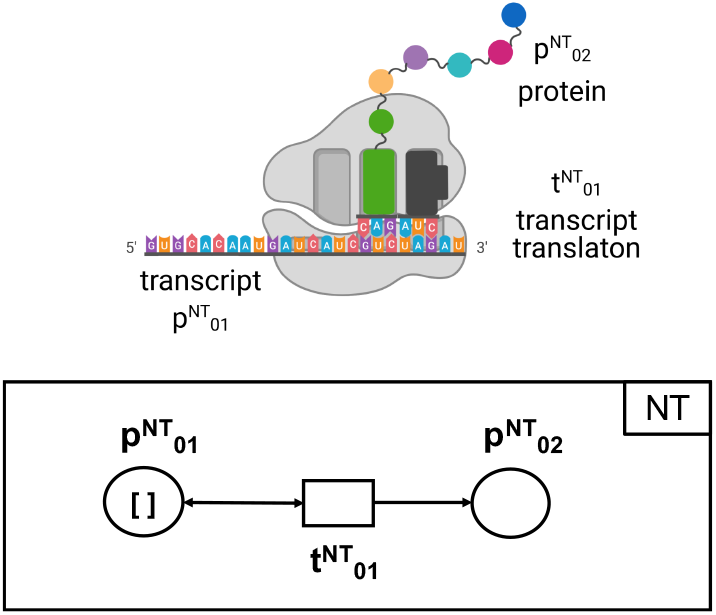
Translation. It models the translation of an RNA transcript into a protein product. This module includes a net-token place for the RNA molecules to be translated 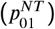 and one for the translation products 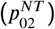, while 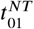models the translation process.

**Figure 31:**
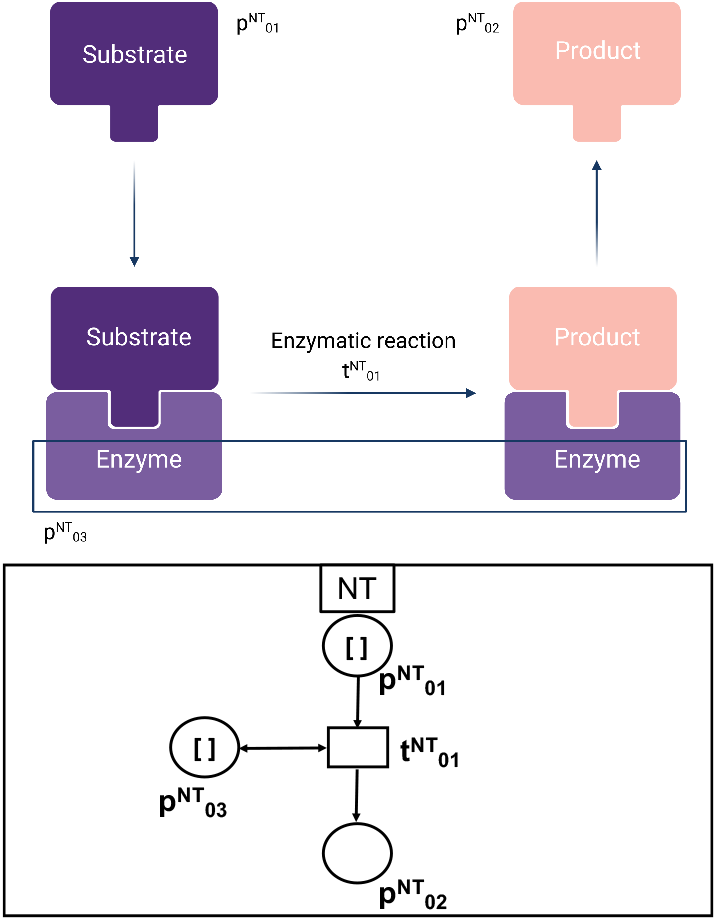
Enzymatic reaction. It models the enzymatic catalysis transforming substrates into products. This module includes a net-token place for the enzymatic reaction substrates 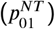, one for the products 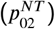, and one for the enzyme to catalyze the reaction 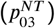, while 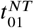 models the reaction process, which depends on the presence of the enzyme.

**Figure 32:**
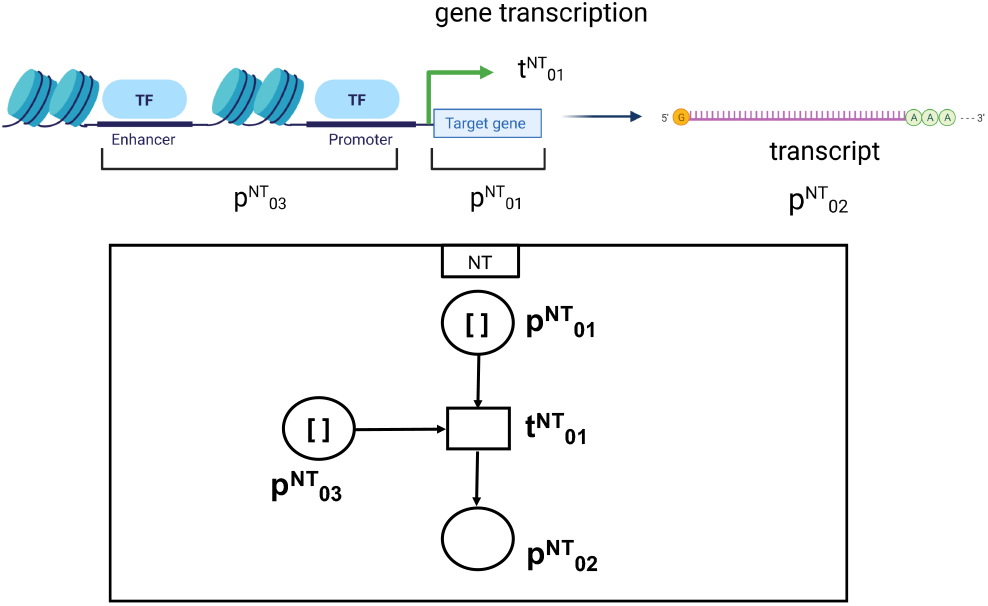
Activating gene regulation. It models the positive regulation of a transcription process. This module includes a net-token place for the gene to be transcribed 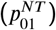, one for the transcription products 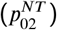, and one for the regulatory signal (for example, a transcription factor) activating transcription 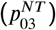, while 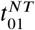 models the transcription process, which depends on the presence of the activating signal.

**Figure 33:**
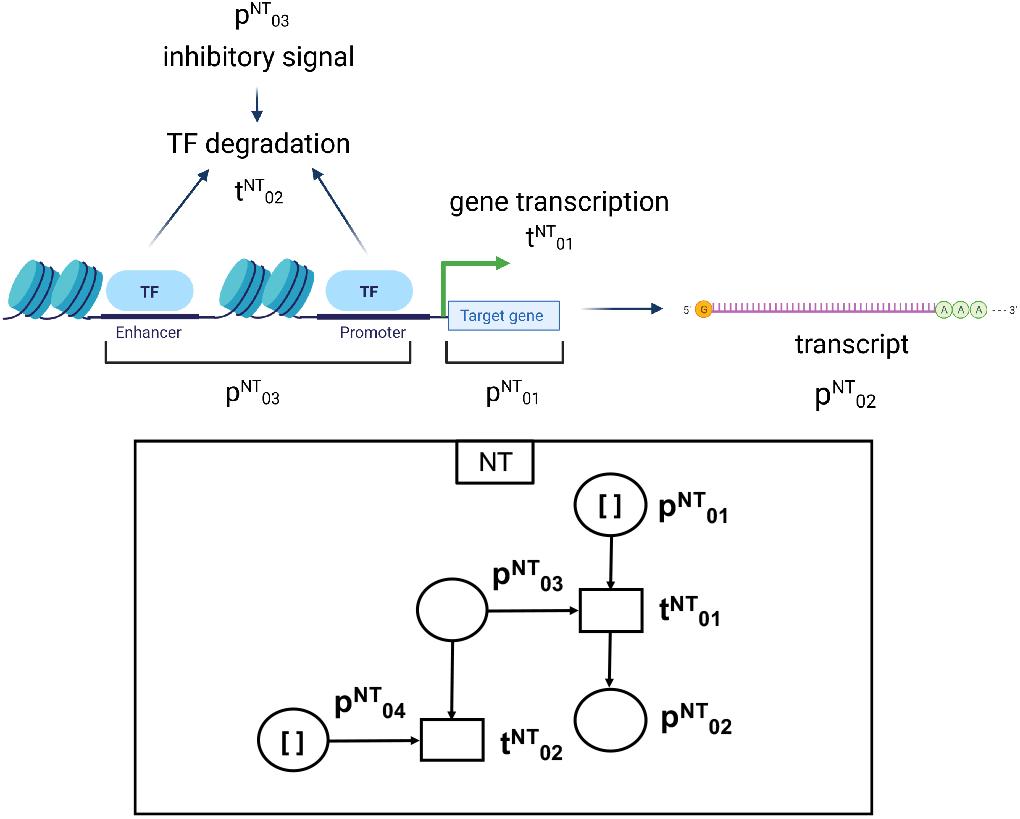
Inhibiting gene regulation. It models negative regulation of a transcription process. This module includes net-token places for the gene to be transcribed 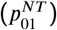, the transcription products 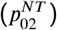, the regulatory signal activating transcription 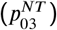, and the signal inducing the degradation of the activating signal 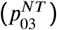. While 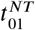models the activation-dependent transcription process, 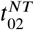 models the degradation of the activating signal, that is induced by the inhibitory signal, resulting in the inhibition of the transcription process.

